# Beyond BOLD: Evidence for diffusion fMRI contrast in the human brain distinct from neurovascular response

**DOI:** 10.1101/2021.05.16.444253

**Authors:** Wiktor Olszowy, Yujian Diao, Ileana O. Jelescu

## Abstract

Functional Magnetic Resonance Imaging (fMRI) is an essential method to measure brain activity non-invasively. While fMRI almost systematically relies on the blood oxygenation level-dependent (BOLD) contrast, there is an increasing interest in alternative methods that would not rely on neurovascular coupling. A promising but controversial such alternative is diffusion fMRI (dfMRI), which relies instead on dynamic fluctuations in apparent diffusion coefficient (ADC) due to microstructural changes underlying neuronal activity, i.e. neuromorphological coupling. However, it is unclear whether genuine dfMRI contrast, distinct from BOLD contamination, can be detected in the human brain in physiological conditions. Here, we present the first dfMRI study in humans attempting to minimize BOLD contamination sources and comparing functional responses at two field strengths (3T and 7T). Our study benefits from unprecedented high spatio-temporal resolution, harnesses novel denoising strategies and examines characteristics of not only task but also resting-state dfMRI. We report task-induced decrease in ADC with temporal and spatial features distinct from the BOLD response and yielding more specific activation maps. Furthermore, we report dfMRI resting-state functional connectivity which, compared to its BOLD counterpart, is essentially free from physiological artifacts and preserves positive correlations but preferentially suppresses anti-correlations, which are likely of vascular origin. A careful acquisition and processing design thus enable the detection of genuine dfMRI contrast on clinical MRI systems. As opposed to BOLD, diffusion functional contrast could be particularly well suited for low-field MRI.

## Introduction

Functional MRI (fMRI) has become an invaluable tool for neuroscience and is nearly omnipresent in studies of healthy and diseased brain. The most widely used fMRI contrast originates from changes in venous blood oxygenation underlying neuronal activity, also known as BOLD (blood oxygenation level-dependent) signal (1), to the extent that “*fMRI*” and “*BOLD fMRI*” have become near-synonyms. However, within the BOLD mechanism also lies the major drawback of this contrast: it relies on neurovascular coupling instead of direct neuronal activity. This additional convolution layer has several dramatic implications (2): loss in spatial and temporal specificity, since neuronal activity is convolved with a slow hemodynamic response function, and strong BOLD signal stems not from the locus of neuronal firing, but from nearby venules and veins (3), a challenging distinction between neuronal and vascular alterations, of relevance to healthy aging or pathology such as tumors and multiple sclerosis (4–6) and low sensitivity in white matter (7, 8), where vasculature and metabolism are reduced. In addition, BOLD is by design sensitive to fluctuations in local magnetic field homogeneity, which can have numerous sources other than blood oxygenation changes. In other words, BOLD fMRI is also notoriously corrupted by physiological noise. Although a battery of processing tools has been developed to correct for these effects, BOLD functional connectivity studies are still mainly used to inform about population differences and exhibit very limited diagnostic performance at the single-subject level (9).

**Diffusion functional MRI (dfMRI)** was proposed as a functional contrast which does not rely on neurovascular coupling (10) but instead on the sensitivity of the apparent diffusion coefficient (ADC) of water to dynamic microstructure fluctuations that typically occur in neurons and astrocytes (including their processes) as a result of neuronal activity (11–13). Indeed, diffusion MRI (dMRI) encodes the mean displacement of water molecules, which is on the order of a few microns and is largely governed by the layout of cellular membranes, compartment sizes and other features of tissue microarchitecture. Since the initial work by Le Bihan *et al*. (14), a body of conflicting evidence has been published. Several studies have reported improved temporal and spatial specificity of the dfMRI signal compared to BOLD (14–16) and specificity to neuronal activity rather than vascular response (17–19). Other works have argued on the contrary that the so-called dfMRI signal is based on residual vascular rather than neuronal contrast and that the method is otherwise not sensitive enough to detect physiological levels of brain activity on an individual basis (20–23). The conflicting literature suggests that achieving genuine dfMRI contrast constitutes a true methodological challenge (**Figure 1**).

**Figure 1.**
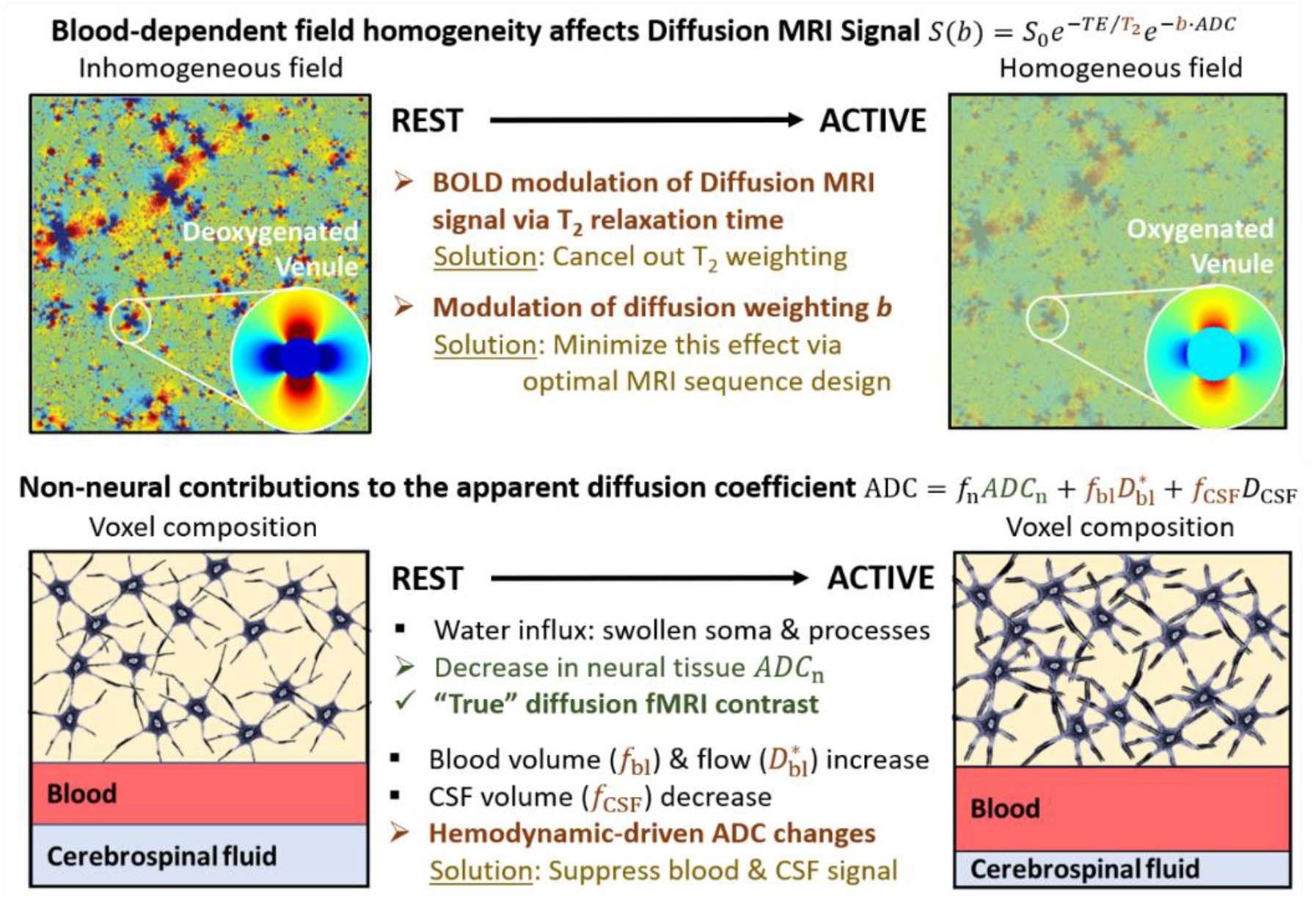
Vascular confounds in dfMRI. Pathways for wanted dfMRI contrast (green font) and for unwanted “contamination” of dfMRI neuronal contrast by the same changes in blood oxygenation underlying the BOLD contrast (dark red font) and solutions to minimize them. Top panels adjusted from Miller and Jezzard, 2008 (Copyright © 2008 Wiley-Liss, Inc.).

Briefly, there are three main mechanisms by which the hemodynamic response to neuronal activity can interfere with the dfMRI contrast, with careful solutions to mitigate them (**Figure 1**, **Table 1**). First, the diffusion-weighted MRI signal is, by design of the spin-echo sequence, also *T*_2_-weighted (or *T*_2_*-weighted, for gradient-echo sequences). Changes in tissue *T*_2_(*) due to blood oxygenation differences between rest and active states are the very mechanism behind the BOLD contrast. Thus, a plain diffusion-weighted signal is also heavily affected by the same mechanism. This confound was present, among others, in (10, 22, 20, 15). Therefore, the ADC time-course should be considered instead – estimated, for instance, from consecutive measurements with two different *b*-values, assuming *T*_2_ to be constant during these two measurements. Second, blood oxygenation dependent field homogeneity implies background field gradients **G**_s_ vary with the hemodynamic response. These gradients can contribute to the effective diffusion-weighting imparted in the sequence, most notably via cross-terms with the diffusion gradients **G**_d_. Contributions from background gradients may have challenged the interpretation in (14, 22, 24). A sequence design that minimizes cross-terms should be preferred, e.g. a twice-refocused spin-echo (25). Third, the ADC at the voxel level is a combination of diffusivities from all voxel constituents: compartments of interest (intra- and extra-neuronal space), but also blood. The hemodynamic response translates into an increase in blood volume and flow (pseudo-diffusion coefficient *D*_bl_*). Blood water contributions to the MRI signal should therefore be eliminated to access changes in tissue ADC only. The inclusion of *b*=0 in the ADC estimation may result in BOLD-like response functions as, for instance, reported in (16, 24). It is noteworthy that techniques to suppress the blood signal by injection of magnetic particles (21) may also introduce an indirect vascular contribution, as they generate susceptibility gradients around vessels that fluctuate with changes in vessel size.

**Table 1.** Explicit mechanisms for vascular contamination of dfMRI contrast, either via the diffusion-weighted signal (Items 1 – 2) or the compartment contributions to the voxel ADC (Items 3 – 4) and technical solutions to address them.

|  | Sources of vascular contamination | Solution |
| --- | --- | --- |
| 1 | Intrinsic $T_2$-weighting in diffusion-weighted signal:<br>$S = M_0 e^{-TE/T_2} e^{-b \cdot ADC}$ | Calculate ADC time-courses from interleaved acquisition of two $b$-values to cancel out $T_2$ weighting:<br>$ADC(t) = \ln\left(\frac{S_2(t + \delta t)}{S_1(t)}\right) \frac{1}{b_1 - b_2}$ |
| 2 | Modulation of effective $b$-value by cross-terms between venous background field gradients $G_s(t)$ and diffusion gradients $G_D$:<br>$b = xG_D^2 + yG_D \cdot G_s(t) + zG_s^2(t)$<br>Coefficients $x, y, z$ are sequence-dependent.<br>Typically $G_D \gg G_s$. | Diffusion gradient waveform design that minimizes cross-terms<br>e.g. twice-refocused spin-echo |
| 3 | Contribution of blood water pool | Operate at $b$-values $\geq 0.2 \text{ ms}/\mu\text{m}^2$ to suppress fast moving spins (blood signal) |
| 4 | CSF volume shrinkage to compensate for blood volume increase | Selectively null CSF signal using inversion module |

Encouragingly, studies of perfused tissues without vasculature have clearly demonstrated that neuronal activity decreases the overall diffusion coefficient (17, 18, 26), although the sensitivity to physiological levels of activity has been questioned (23). New approaches of ultra-high temporal resolution to separate early dfMRI from later BOLD effects have nonetheless singled out genuine dfMRI contrast in the rodent (16, 27).

We adopt a dfMRI study design that minimizes the aforementioned sources of vascular contamination to determine whether genuine diffusion fMRI contrast is detectable in the human brain at the individual level. A bipolar gradient diffusion sequence is used to mitigate cross-terms between diffusion and susceptibility gradients, and apparent diffusion coefficient (ADC) time-courses are computed using *b*-values ≥ 0.2 ms/μm^2^ to suppress *T*_2_-weighting and blood pool contributions. We explore different approaches to assessing residual vascular contributions by comparing the dfMRI signal properties at two field strengths, 3T and 7T. From a biophysical standpoint, the dfMRI response should be field-independent while the BOLD magnitude increases with field strength. Finally, we evaluate dfMRI properties not only in response to a task but also, for the first time, in terms of resting-state functional connectivity (RS-FC) compared to *T*_2_-BOLD. The notoriously low sensitivity of dfMRI is balanced by recent improvements in spatio-temporal resolution of the acquisition (e.g. stronger gradients on clinical systems and multi-slice acceleration) and in image pre-processing, in particular Marchenko-Pastur PCA (MP-PCA) denoising (28–30) to boost the temporal signal-to-noise ratio (tSNR).

## Results

Data were acquired on Siemens Magnetom 7T and Prisma 3T scanners, both equipped with 80 mT/m gradients, on 22 subjects (7 males, age 25 ± 5). Four scanning protocols were compared: (1) SE-EPI yielding *T*_2_-BOLD contrast, (2) DW-TRSE-EPI with pairs of *b*-values 0.2 and 1 ms/μm^2^ (our *optimized* dfMRI protocol), (3) DW-TRSE-EPI but with *b*-values 0 and 1, and (4) DW-SE-EPI (monopolar gradient pulses) with *b*-values 0.2 and 1. The pair of *b*-values=0.2/1 was chosen to satisfy three criteria: suppress direct vascular signal (*b* ≥ 0.2 ms/μm^2^) (21), provide a sufficient range for ADC estimation and preserve sufficient SNR in individual diffusion-weighted images.

**Table S1** specifies the numbers of datasets retained for the analyses (N=64 total for task, and N=55 total for resting-state, all protocols combined). Six datasets were excluded from the analyses due to substantial outliers across timecourses. Two resting-state datasets were excluded due to inconsistent positioning of the imaging slab. Data from one subject was excluded entirely because of poor task compliance. Finally, five datasets were excluded because of image artifacts.

The relatively short TR of 1 second, chosen in the interest of high temporal resolution, yielded limited steady-state signal and SNR. However, the MP-PCA denoising procedure nearly doubled the tSNR (**Fig. S1**).

For task datasets, our main goal was to compare response functions across the modalities and field strengths. For resting-state datasets, our main goal was to compare functional connectivity matrices, also both across the modalities and field strengths.

### Task fMRI

The task consisted in viewing a flashing checkerboard (8 Hz) and concurrently finger-tapping with both hands for 12s following 18s of rest. **Figure 2** shows group-level activation maps resulting from Protocols (1) and (2) at each field strength. Activation in the visual and motor cortices was observed, both with SE BOLD and with dfMRI, at 3T and 7T. While the sensitivity of dfMRI is expectedly lower, its activation patterns point towards improved specificity: the voxels consistently active at the group-level delineate cortical ribbons in the visual area, interhemispheric supplementary motor area and bilateral motor/somatosensory cortices.

**Figure 2.**
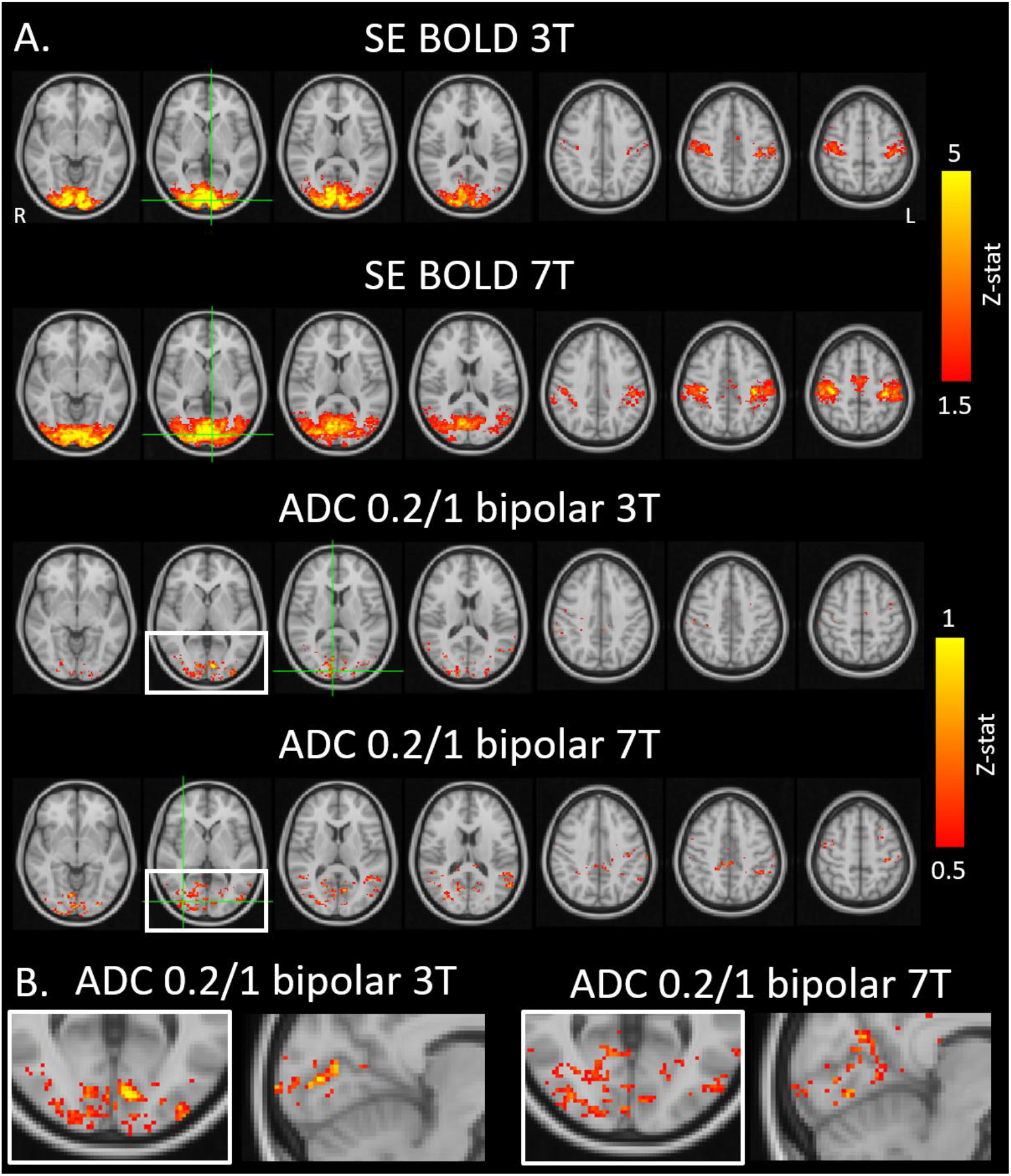
Whole-brain group-averaged z-statistic maps for SE BOLD and dfMRI, at 3T and 7T. To find task-induced activation without making assumptions about the shape of the response functions, a Finite Impulse Response (FIR) set was used. Significance was assessed with F-tests, which were converted to z-statistics. **A**: Activation both in the visual and motor cortices was observed, both with SE BOLD and with dfMRI, at both fields. Green crosshairs mark the cluster peak. **B**: While the sensitivity of dfMRI is expectedly lower, its activation patterns point towards improved specificity: the voxels consistently active at the group-level delineate cortical ribbons in the visual area, interhemispheric supplementary motor area and bilateral motor/somatosensory cortices.

Analysis of the group-averaged response functions revealed, as expected, a positive SE BOLD response with an amplitude larger at 7T than at 3T (2% vs. 1% in the visual cortex), as well as a delayed BOLD onset with respect to stimulus and even more delayed return to baseline after stimulus end **(Figure 3A**). It should be noted that these BOLD amplitudes are expectedly lower for *T*_2_-BOLD than for the more familiar *T*_2_*-BOLD. Our optimized dfMRI protocol revealed a task-induced ADC decrease by around 1% **(Figure 3B**). Unlike the delayed BOLD response, the ADC decrease onset was concomitant with the stimulus. At 3T, the decrease was gradual and peaked around 8 – 9 s into the stimulus before increasing back to baseline. The return to baseline was complete by the end of the paradigm block in the motor cortex and slower in the visual cortex, up to 8 seconds from the stimulus end. At 7T, the response function in the motor cortex was qualitatively similar to 3T but with a larger amplitude. In the visual cortex, the response was non-monotonic with an initial decrease in ADC, similarly to 3T, followed by a very rapid return to baseline and post-stimulus over-shoot in ADC. These main time-courses are overlaid in **Figure 4** for better comparison. Illustrative examples of individual responses to task for Protocol (2) (ADC *b*=0.2/1 bipolar) are shown in **Fig. S2**.

**Figure 3.**
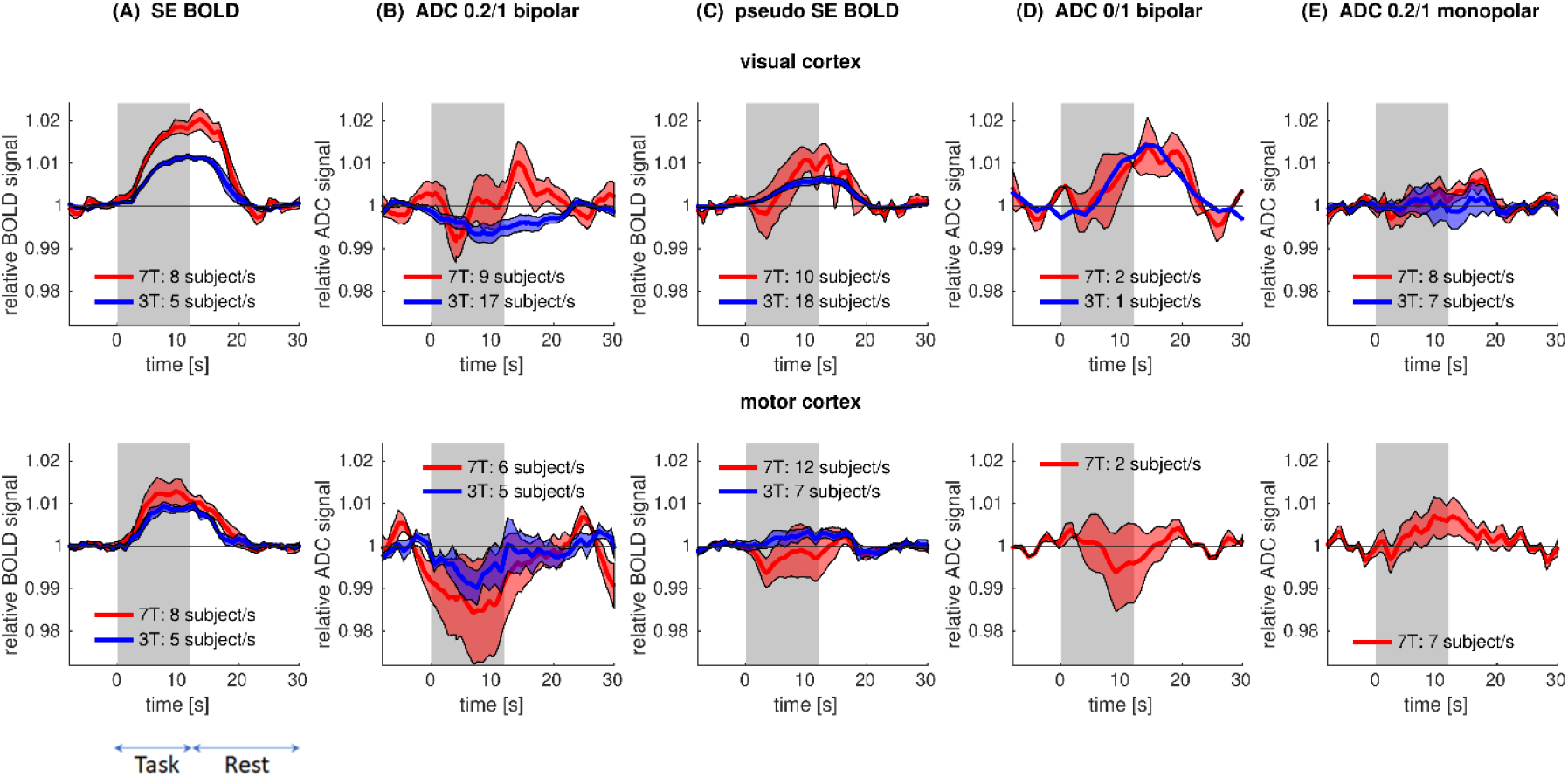
Group-averaged response functions for SE BOLD (**A**), dfMRI from b=0.2/1 pairs bipolar (**B**), pseudo SE BOLD (**C**), dfMRI from b=0/1 pairs bipolar (**D**), and dfMRI from b=0.2/1 pairs monopolar (**E**), at 3T and 7T. Shaded area: SEM. **A:** The positive SE BOLD response displayed an amplitude larger at 7T than at 3T (2% vs. 1% in the visual cortex), as well as a delayed BOLD onset with respect to stimulus and even more delayed return to baseline after stimulus end. **B:** The optimized dfMRI protocol revealed a task-induced ADC decrease by around 1%. The sign and temporal characteristics of the response were markedly different from BOLD, as detailed in **Fig. 4**. **C – E:** The qualitative trends of the response functions were similar to BOLD, pointing to the importance of each step in our optimized dfMRI (Column B) to minimize vascular contributions. For example, a pseudo T_2_-BOLD with small diffusion-weighting – likely suppressing the contribution from blood spins – still revealed BOLD-like response, particularly in the visual cortex (Column C). Datasets for b=0/1 ADC show an increase in ADC during task in the visual cortex, similarly to BOLD, but a decrease in the motor cortex (Column D). Finally, the protocol with monopolar diffusion gradient pulses resulted in a nearly flat response in the visual cortex and an increase in ADC in the motor cortex (Column E).

**Figure 4.**
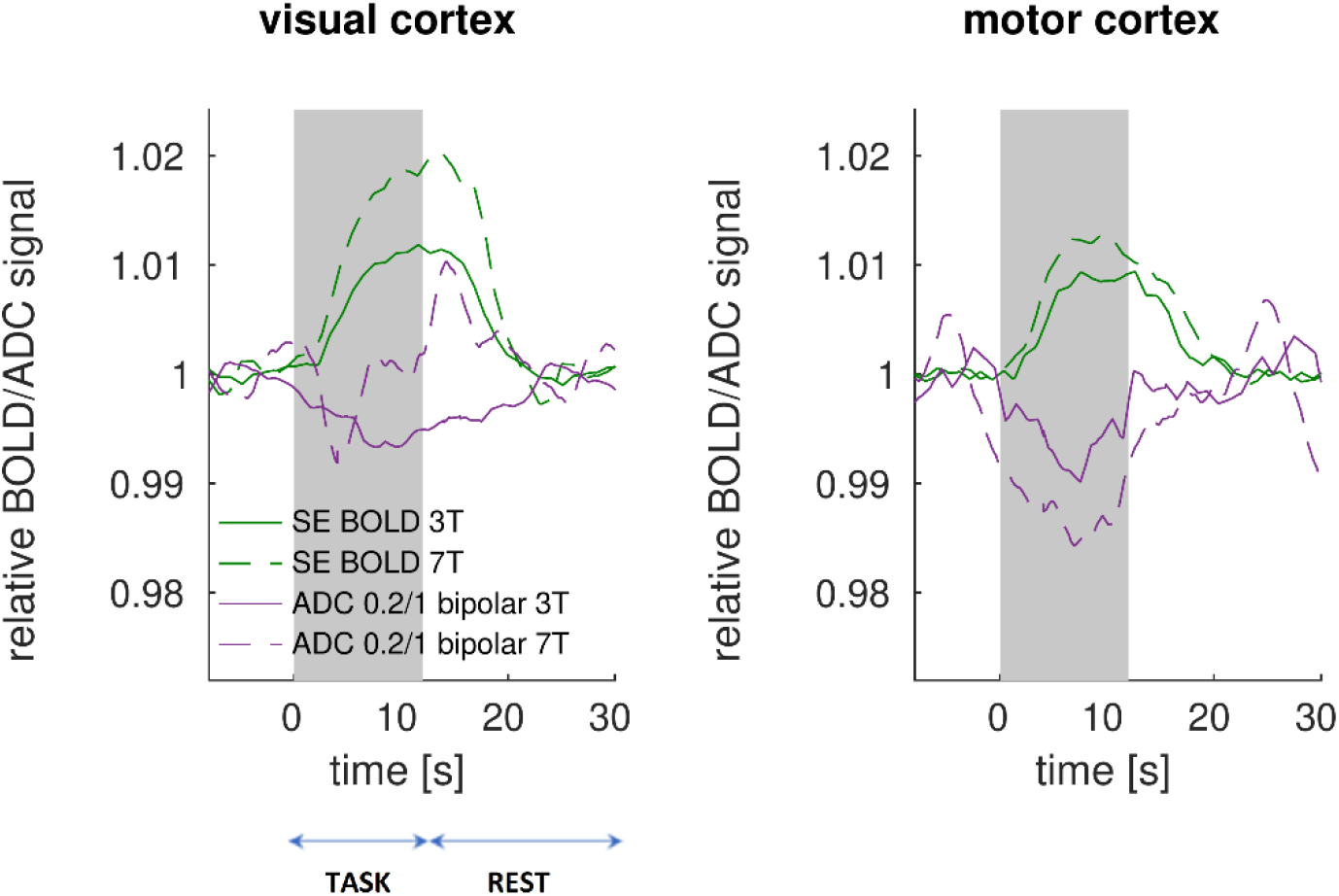
Group average response functions overlaid for SE BOLD and dfMRI at 3T and 7T in the visual and motor cortices. Unlike the delayed BOLD response, the ADC decrease (dfMRI) was gradual but immediate from stimulus onset. At 3T, dfMRI response peaked around 8 – 9 s into the paradigm before increasing back to baseline. The change in ADC directionality occurred while the BOLD response was sustained at a plateau. At 7T, the dfMRI response function in the motor cortex was qualitatively similar to 3T but with a larger amplitude. In the visual cortex, the response was non-monotonic with an initial decrease in ADC, similarly to 3T, followed by a very rapid return to baseline and post-stimulus over-shoot in ADC.

For the other acquisition protocols **(Figure 3, C – E**), the qualitative trends of the response functions were similar to BOLD, pointing to the importance of each step in our optimized dfMRI (Column B) to minimize vascular contributions. For example, *b*=0.2 time-courses analyzed independently as a pseudo *T*_2_-BOLD with small diffusion-weighting – likely suppressing the contribution from blood spins – still revealed BOLD-like response due to *T*_2_-weighting, particularly clear in the visual cortex (Column C). Few datasets are available for *b*=0/1 ADC, but they show nonetheless an increase in ADC during task in the visual cortex, similarly to BOLD, but a decrease in the motor cortex (Column D). Finally, the protocol with monopolar diffusion gradient pulses resulted in a nearly flat response in the visual cortex and an increase in ADC in the motor cortex (Column E).

### Resting-state functional connectivity

Exemplary signal and noise ICA components are shown in **Fig. S3**. Group averages of FC matrices following manual ICA cleaning and GSR were dominated by strong positive correlations, which agreed remarkably well between *T*_2_-BOLD and dfMRI methods, at both field strengths (**Figure 5**). However, our optimized dfMRI protocol attenuated anti-correlations preferentially compared to *T*_2_-BOLD, particularly at 3T. Indeed, the slope of the linear regression of dfMRI to *T*_2_-BOLD positive correlations was 0.93 at 3T and 0.92 at 7T, while for *T*_2_-BOLD negative correlations, the slope dropped to 0.28 at 3T and 0.51 at 7T, respectively. *t*-tests of FC strength between the two protocols showed that the majority of edges with significantly different connectivity had either a low or negative correlation in BOLD, which was further attenuated in dfMRI (**Table S2**).

**Figure 5.**
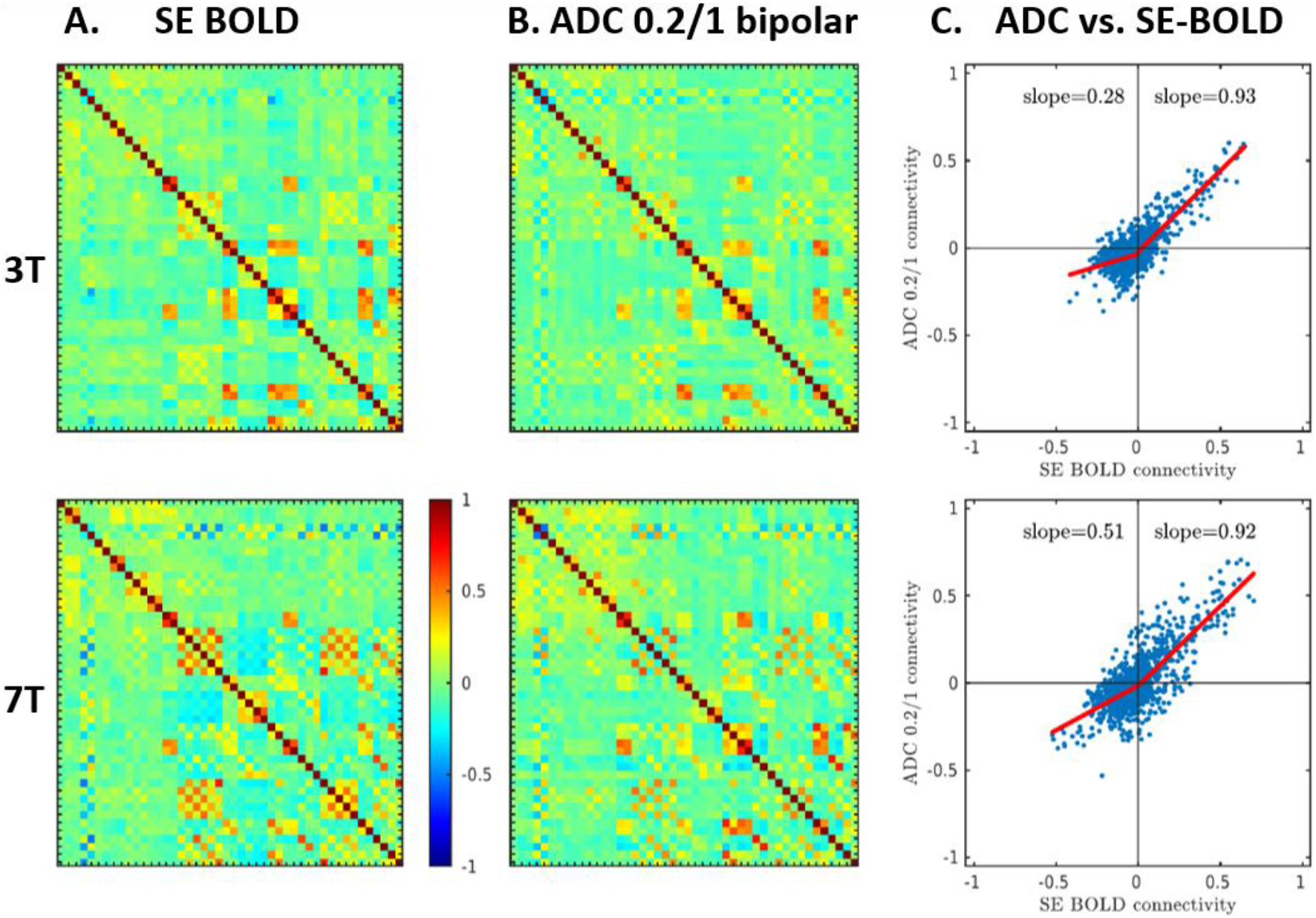
Resting-state analysis following manual ICA cleaning and GSR. Group averages of FC matrices from SE BOLD (**A**) and dfMRI (ADC b=0.2/1 bipolar) (**B**) at 3T and 7T. **C**: Correlation of FC strength derived from SE BOLD vs. dfMRI, assessed separately for positive and negative edges of BOLD fMRI. The positive edges in BOLD fMRI agree exceptionally well with dfMRI FC (slope of the linear regression ^~^1 at both field strengths). However, BOLD negative edges (anti-correlations) are largely suppressed in dfMRI, particularly at the lower field.

Remarkably, FC patterns were very similar between our dfMRI protocol and the blood-attenuated pseudo SE BOLD (**Fig. S4**). As in dfMRI, the anti-correlations were reduced in pseudo SE BOLD and both positive and negative correlations agreed between the two methods. It should be underlined that the FC comparison is all the more powerful between these protocols as the underlying data were acquired during the same run, and thus for identical brain activity and physiological conditions in the volunteer.

Finally, the comparison of how various processing steps impact functional connectivity as estimated from the different protocols (SE BOLD, pseudo SE BOLD and ADC 0.2/1 bipolar) suggested that ICA cleaning had a marked impact on SE BOLD connectivity – by removing unwanted physiological artifacts (65% of ICA components), but less impact on ADC-derived connectivity (33% of ICA components were classified as artifacts) (**Fig. S5**). By intrinsically removing contributions from large field fluctuations to functional contrast, such as those of physiological origin, ADC-dfMRI also has a valuable attribute of yielding resting-state data that is “*clean*” by design.

## Discussion

This is the first dfMRI study attempting to minimize BOLD contamination sources and compare functional responses at two field strengths. It is also the first study in humans that examines not only task but also resting-state dfMRI features. Our study further benefits from unprecedented high spatial and temporal resolution of 2 – 2.5 mm isotropic and 1 – 2 s. Most of the previous human dfMRI studies employed lower spatial resolution, up to 4 mm isotropic (14), and lower temporal resolution, up to 8 s (24), which obscured the interpretation of spatial specificity and temporal characteristics of the functional response.

### Task fMRI

The optimized dfMRI protocol was the only one to yield a consistent decrease in ADC during task, as expected from biophysical mechanisms and previously reported in experiments on tissue samples without vasculature (17, 18, 26). All other “*dfMRI*” protocols yielded BOLD-like response functions (from *T*_2_-weighting or direct blood pool contributions), with a directionality not fully consistent across ROIs or field strengths. This inconsistency (e.g. for ADC time-course from *b*=0/1, between the visual and motor cortices) could be related to spatially varying SNR levels, field homogeneity, magnitude of the neuronal response and of a competing BOLD response. The ADC time-course from *b*=0.2/1 pairs with monopolar gradient pulses showed a weak, nearly flat response at 3T, potentially resulting from competing dfMRI and BOLD mechanisms balancing out the net effect.

A close comparison between *T*_2_-BOLD and optimized dfMRI characteristics suggests different contrast mechanisms between the two. Indeed, the BOLD response was delayed with respect to the stimulus onset and, once established, the plateau lasted at least until the stimulus end (motor cortex) or beyond (visual cortex) before gradually decreasing to baseline. Return to baseline was reached within eight seconds from the stimulus end. The differences in BOLD response between cortical areas may be due to a difference in stimulus strength and persistency as well as in hemodynamic response (4, 31). In contrast, for both ROIs and at both field strengths, the ADC decrease started upon stimulus onset and time to trough was 8 – 9 seconds into the stimulus. In the visual cortex at 3T, the ADC started increasing back towards baseline at the end of the stimulus, while the BOLD plateau was maintained beyond. In the visual cortex at 7T, there was a more rapid return of ADC to baseline and a positive over-shoot after the stimulus period. The first component might reflect rapid microstructural changes, and the latter slower but notable BOLD contamination, in full agreement with a recent dfMRI study at 9.4T in rats (27).

The difference in dfMRI qualitative trend between 3T and 7T suggests vascular contributions may be more challenging to mitigate at higher fields due to stronger susceptibility, despite a careful sequence design. In the motor cortex, the dfMRI responses were qualitatively more consistent between 3T and 7T, with a gradual decrease initiated without delay from stimulus onset and 8-second time to trough, followed by a gradual return to baseline after stimulus end. BOLD effects may be brain-region dependent and less pronounced in the motor vs. the visual cortex (4, 31). It is worth noting, for instance, that BOLD amplitude difference between 7T and 3T was very pronounced in the visual cortex, but not so marked in the motor cortex. However, the magnitude of the dfMRI effect (via the ADC decrease) does appear to be field-dependent also in the motor cortex, whereby vascular contributions to dfMRI cannot be ruled out in this ROI at 7T.

The relatively smooth and persistent ADC decrease during task is consistent with previous dfMRI studies (14, 26) and also with expected gradual changes in the overall volume of neuronal, glial and extracellular compartments upon sustained firing, as highlighted by the electrodiffusive neuron-extracellular-glia model (32).

We report an amplitude of ADC decrease on the order of 1% in physiological conditions of sensory and motor stimulation. This magnitude of the effect is weak but nonetheless detectable, and sensible when compared to preclinical dfMRI studies using chemical or electrical stimulation above physiological thresholds, and which reported ADC drops of 6 – 50% (17, 26). It is also important to note that our estimated response functions were somewhat noisier than in some of the previous literature, where the time-courses were filtered – either with a median filter (27) or with a moving average filter (14, 24). We preferred not to use a filter, as a filter could have hidden subtle ADC dynamics.

As for the temporal characteristics, the spatial distribution of activated voxels was also very different between BOLD and dfMRI. As expected, the smaller dfMRI effect size resulted in a more limited activation pattern and weaker statistics at the group level. Nevertheless, the activated voxels consistently followed cortical ribbons in the active areas, yielding more specific information than large BOLD clusters. While cortical layer-specific activation patterns can be achieved with BOLD or VASO fMRI at ultra-high fields while scanning at very high spatial resolution, particularly strikingly with laminar fMRI (33), with dfMRI similar patterns were accessible using much less demanding settings: a 3T field strength and a spatial resolution of 2.5 mm (which corresponds to the average human cortical thickness). Furthermore, the BOLD cluster peaks did not coincide spatially with dfMRI peaks, suggesting dfMRI contrast is distinct from a residual strong BOLD contamination. We note that our implementation of BOLD corresponded to *T*_2_-BOLD, which has improved sensitivity towards capillaries than the more common *T*_2_*-BOLD, where the largest signals stem from the draining veins. DfMRI therefore shows improved specificity even over a BOLD technique refined towards capillaries.

The analysis of the spatial location of active voxels with either method also highlighted a larger proportion of active voxels in major white matter tracts – as labelled in the Johns Hopkins University atlas (34) – with dfMRI (5 – 7%) vs. SE BOLD (1%). It is well-established that BOLD sensitivity in white matter is much reduced due to lower blood volume and flow as well as energy requirements compared to gray matter. As a result, most resting-state and some task BOLD fMRI studies still use the white matter signal as a nuisance regressor. Although white matter BOLD is regaining interest (7, 8) – possibly thanks to higher field strengths and spatial resolution boosting sensitivity and specificity – the positive findings in this field remain limited. DfMRI has been singled out as a promising technique to detect white matter activity (35), and a decrease in ADC upon sustained stimulation was measured in *ex vivo* nerve sample or mouse optic nerve (26, 36, 37). Our results support the stronger potential of dfMRI vs. BOLD to detect white matter activity. This potential requires, however, further investigation given the more limited sensitivity of dfMRI overall.

### Resting-state functional connectivity

Based on resting-state data that underwent ICA cleaning and GSR, or GSR alone, we found primarily reduced anti-correlations and maintained positive correlations in dfMRI RS-FC vs. BOLD, which supports two hypotheses.

First, that GSR does not result in a plain mathematical demeaning of FC matrices but contributes to highlighting genuine (anti-)correlations across the brain, should they exist. This finding is in agreement with recent studies – both in humans and rodents – showing that ICA cleaning + GSR also improves the differentiation between populations based on resting-state patterns compared to ICA cleaning alone (29, 38). While GSR is known to mathematically favor anti-correlations (39), there is also evidence of genuine anti-correlations in BOLD resting-state functional connectivity, which are reportedly related to certain brain networks being specifically inactivated while other networks are active, for accrued efficiency (40), e.g. default-mode vs. executive control networks (41, 42). Thus, they could have the same vascular origin as negative BOLD, related to arteriolar vasoconstriction and reduced oxygenation in areas of suppressed neuronal firing (43, 44).

Second, and most importantly for our purposes, this difference in RS-FC patterns between BOLD and dfMRI suggests that there is no twin of negative BOLD in dfMRI and that the sources of dfMRI signal are distinct from those of BOLD and largely free of vascular contributions, in agreement with previous findings in the rat brain (45, 46). Indeed, from the perspective of neuron microstructural fluctuations, inhibitory activity leading to membrane hyperpolarization is not expected to alter the cell volume and/or morphology substantially relative to baseline – i.e. the change in membrane potential between resting and hyperpolarized states is small compared to the dramatic change in membrane potential incurred during firing (47).

We note that the pseudo SE BOLD data (based on *b*=0.2 ms/μm^2^ time-courses) yielded RS-FC with attenuated anti-correlations and that agreed remarkably well with dfMRI RS-FC coefficients. Therefore, it would appear that BOLD anti-correlations could be driven by signal contributions from the blood pool, which are already largely attenuated using mild diffusion-weighting.

Finally, while ICA cleaning had a large impact on BOLD RS-FC, the effect was lesser on dfMRI RS-FC. This is consistent with the fact that ICA components typically identified as artifacts are largely of physiological origin, be it variations in heart rate, breathing rate, or motion that affect the field homogeneity and contribute to non-BOLD *T*_2_(*) fluctuations. From this perspective, dfMRI time-courses can be considered “*self-cleaned*”, which represents a major advantage in limiting pre-processing steps and reducing within- and between-subject confounding variability.

### Limitations

In the interest of temporal resolution, this study used a short TR of 1 s, atypical for multi-slice SE-EPI and which only allowed limited *T*_1_ recovery – especially at 7T – resulting in limited SNR. Furthermore, the *B*_1_ transmit field inhomogeneity is notoriously pronounced at high field, further causing signal loss for a spin-echo and all the more for a double spin-echo sequence design. The relatively low SNR was, however, mitigated by the MP-PCA denoising approach.

The relatively high temporal resolution also prevented whole-brain coverage. Thus, the analysis of dfMRI responses was confined to a slab covering the areas anticipated as most relevant for either task or resting-state. However, the achievable brain coverage was doubled using multiband acceleration of factor 2, with some further penalty in SNR. Additional brain regions may be of interest, particularly given the spatial mismatch between BOLD and dfMRI cluster peaks (i.e. voxels with the strongest activation in either case) and between RS-FC patterns. Overall, our acquisition protocol achieved a sensible trade-off between temporal resolution, spatial resolution, SNR and brain coverage. Still, ongoing efforts in acquisition strategies, acceleration and efficiency may allow to further optimize these parameters concomitantly.

Both the task and resting-state results showed that the dfMRI signal at 7T shared combined features of dfMRI at 3T and BOLD. This suggests that despite a careful experimental design, vascular contributions to dfMRI cannot be suppressed entirely, especially at higher field strengths, where susceptibility effects are more pronounced.

All of the above limitations point towards dfMRI being perhaps better suited for lower field strengths. Interestingly, now in the MRI community, there is an increasing interest in low-field systems (48, 49). While lower fields are detrimental to SNR and CNR in BOLD fMRI, and laminar fMRI in particular, for dfMRI, low-fields may yield acceptable SNR (via shorter *T*_1_ and more homogeneous *B*_1_), larger brain coverage (via reduced radio-frequency power deposition and heating) and cleaner, more specific contrast (via reduced BOLD contributions).

One potential indirect BOLD contribution that our protocol did not account for is a decrease in CSF volume as a compensation mechanism for an increase in blood volume during the hemodynamic response (50, 51). However, diffusion-weighting is expected to also attenuate the contribution from the fast-diffusing CSF compartment quite substantially, reducing the impact of this effect. Future dfMRI protocols could nonetheless include an initial CSF-nulling inversion pulse (52) to address this confound directly.

Task dfMRI sensitivity could be improved if the shape of the response could be assumed to be known a priori – like in the BOLD fMRI studies, where usually the canonical hemodynamic response function (31) is used. We used an FIR model with seven basis functions, which lowered the sensitivity compared to a scenario where the shape of the response was known a priori and the GLM involved one basis function only. While the non-ADC-dfMRI response shape is well-established (53), more work is needed to investigate if a “*canonical*” ADC-dfMRI response function could be used. Another way of increasing dfMRI sensitivity could be to calculate ADC using a sliding window of two time-points. For example, volumes acquired for a sequence of interleaved *b*-values (0.2, 1, 0.2, 1,…) could be used to calculate ADC from *b*-values (0.2/1, 1/0.2, 0.2/1,…). This way, the temporal resolution of ADC time-courses would be doubled for the GLM analysis, and with appropriate pre-whitening, the temporal autocorrelation would be accounted for. We also note the pre-whitening algorithms in other software packages (AFNI or SPM with FAST option) may further improve the spatial specificity of task dfMRI (54).

### Conclusions

Taken together, our task and resting-state results support the existence and detectability of genuine dfMRI contrast distinct from BOLD mechanisms, made possible by a careful study design to minimize vascular contributions. DfMRI enables a more specific detection of activation in response to task and resting-state functional connectivity mapping directly free from physiological artifacts. Finally, dfMRI is ideally suited for low or intermediate magnetic field systems. This will become an important asset in the context of increasing interest in low-field systems, and provide a unique functional MRI technique in a context where BOLD may lack sensitivity.

## Materials and Methods

### Experimental

The study was examined and approved by the ethics committee of the canton of Vaud (CER-Vaud). Twenty-two subjects (7 males, age 25 ± 5) were scanned. All subjects gave informed written consent after the experimental procedures were explained and prior to enrollment. Data were acquired on Siemens Magnetom 7T and Prisma 3T scanners, equipped with 80 mT/m gradients, using a 32-channel (7T) or a 64-channel (3T) receiver head coil. Four scanning protocols were used: (1) SE-EPI yielding *T*_2_-BOLD contrast, (2) DW-TRSE-EPI with pairs of *b*-values 0.2 and 1 ms/μm^2^, (3) DW-TRSE-EPI but with *b*-values 0 and 1, and (4) DW-SE-EPI (monopolar gradient pulses) with *b*-values 0.2 and 1. Sequences were obtained from the Center for Magnetic Resonance Research of the University of Minnesota (https://www.cmrr.umn.edu/multiband/), (55). All scanning was performed with GRAPPA=2 and Multiband=2 acceleration. Sequence parameters and the numbers of datasets are collected in **Table S1** for each protocol, field strength and scan type (task or resting-state). While most of the task datasets were acquired at an isotropic resolution of 2.5 mm, the first datasets had an isotropic resolution of 2 mm. The voxel size was increased in order to improve the SNR. All resting-state datasets were acquired at an isotropic resolution of 2 mm. The scan time for each functional run was approximately 10 minutes.

For all resting-state datasets and most of task dfMRI datasets (13 out of 22 subjects), the two *b*-values were acquired in an alternating fashion within the same functional run. The temporal resolution of the ADC map was thus twice the repetition time (TR), and half the temporal resolution of SE-BOLD. In a few dfMRI task datasets, the two *b*-values were acquired in separate runs to keep the temporal resolution consistent with SE-BOLD. ADC was then calculated from the two separate acquisitions. However, this design was found to be often detrimental in terms of large-scale fluctuations that did not enable the time-courses to be used for an ADC computation, with little benefit/insight brought by higher temporal resolution.

For task datasets, the imaging slab covered both the visual and motor cortices. For resting-state datasets, the imaging slab covered the prefrontal cortex and the posterior cingulate cortex, which are crucial elements of the default mode network (56). The outermost slices were discarded due to inflow contributions. To be able to later correct the susceptibility distortions using *topup* (57), a few EPI volumes were acquired with an opposite phase encoding direction to enable susceptibility distortion correction. An anatomical 1 mm isotropic *T*_1_-weighted image was acquired using an MPRAGE sequence.

For task fMRI, subjects were viewing a flashing checkerboard (8 Hz) and concurrently finger-tapping with both hands for 12s following 18s of rest. This boxcar paradigm was implemented using PsychoPy (58). For the resting-state runs, subjects were told to fixate on the cross in the middle of the screen and to relax.

### Processing

The processing pipeline is sketched in **Fig. S6**.

### Pre-processing

Volume outliers, characterized by signal dropouts either due to scanner instabilities or motion, were automatically detected and replaced with a linearly interpolated signal. A volume was considered an outlier if its linearly detrended average signal over the entire brain was off by more than a pre-specified fraction of the median across all the volumes. This threshold fraction was set at 1% and 3% for 3T and 7T, respectively. It should be noted that after whole-brain averaging, 1 – 3% deviations are much larger than any BOLD fluctuations within this range at the individual voxel level. In most cases, the fraction of outliers did not exceed 5%. If this fraction was above 20%, the entire dataset was excluded from further analyses. For the dfMRI datasets, outlier detection was performed on volume series corresponding to the two *b*-values separately (**Fig. S7**).

Data were then denoised using the MP-PCA method (28, 30) with a sliding window kernel of 5×5×5 voxels. For the diffusion datasets, the denoising procedure was performed on volume series corresponding to the two *b*-values separately. The residuals were inspected for Gaussianity to verify that the denoising procedure did not alter data distribution properties. Corrections were applied for the Gibbs ringing artifact (59) and the susceptibility distortions (57). For the latter, FSL’s tool *topup* was run with settings taken from the UK BioBank study (60). Motion correction was performed with SPM (61, 62). For the diffusion datasets, motion correction was performed on volume series corresponding to the two *b*-values separately. Afterwards, the two series were aligned with each other using ANTs rigid transformation (63, 64).

As the last stage of the pre-processing of the diffusion data, ADC maps were calculated as: ADC = ln(*S*_2_/ *S*_1_)/(*b*_1_ – *b*_2_), where *S_i_* is the signal level measured with diffusion-weighting *b_i_*. The later dfMRI task and resting-state analyses employed the quantitative ADC timeseries. However, the dfMRI volumes acquired at the lower *b*-value (0.2 ms/μm^2^) were also used independently as an approximation for SE BOLD – in the following, these datasets are called “*pseudo SE BOLD*”. These pseudo SE BOLD time-courses had the advantage of being directly comparable to the matching ADC time-courses due to their concomitance in time, with identical physiological conditions and brain activity in the volunteer. No spatial smoothing was used (65).

### Atlas registration

Both task and resting-state analyses were performed in subject’s native space. All the associated registration was done with ANTs (63, 64). Following brain extraction, the subject’s anatomical image was registered to the Montreal Neurological Institute (MNI) template at 1-mm isotropic resolution using symmetric normalization (SyN). The mean functional image was registered to the subject’s anatomical image using rigid transformation. For diffusion data, the mean functional image was calculated for the lower *b*-value volumes only. For the task analysis, the visual and motor cortices in MNI space were defined using the 1.5 mm isotropic Neuromorphometrics atlas (*Neuromorphometrics*, *Inc*.). The visual cortex was the sum of the calcarine cortex, occipital pole, superior occipital gyrus, inferior occipital gyrus, cuneus, lingual gyrus, middle occipital gyrus and occipital fusiform gyrus. The motor cortex was the sum of the precentral gyrus medial segment, precentral gyrus and supplementary motor cortex. All the regions were considered bilaterally. Using the above-obtained transformations, the two MNI-defined regions of interest (ROIs) were brought to subject’s functional spaces. For the resting-state analysis, the available ROIs from the Neuromorphometrics atlas covered by the partial-brain imaging slab were considered. ROIs for which a coverage of ten or fewer voxels was available were omitted.

### Task fMRI analysis

Whole-brain general linear model (GLM) analyses were performed in FSL (66), using the Finite Impulse Response (FIR) method (67) to find activations without making assumptions about the shape of the responses. Both for the SE BOLD and dfMRI analyses, FIR was used with seven basis functions covering 30s of the post-stimulus-onset time. A high-pass filter with a cutoff of 1/100 Hz was applied, both on the data and the model. *F*-tests were run on the FIR basis functions and *F*-statistics were transformed into *z*-statistics. For each fMRI dataset and each of the two ROIs (visual and motor cortices), the response function was estimated as the average signal across the trials and the voxels with *z*-statistic > 3.1. If at least 20 voxels in the ROI had a *z*-statistic > 3.1, the estimated response function was taken to the group analysis. For the calculation of group averages, subject-level estimates were first normalized using the 8 seconds before the stimulus onset as baseline.

### RS fMRI analysis

Pair-wise Pearson correlation coefficients were calculated between the mean time-courses across each ROI, following global signal regression (GSR), manual independent component analysis (ICA) cleaning (29, 68, 69) with high-pass temporal filtering (f > 0.01 Hz) and 40 independent components, or both. For group analyses, resting-state functional connectivity matrices were averaged across subjects. Only ROIs covered by all subjects were considered at the group level.

## Data and code availability

Data are available on OpenNeuro (currently under a grace period): https://openneuro.org/datasets/ds003676/versions/1.0.0. All the processing scripts needed to fully replicate our study are at https://github.com/wiktorolszowy/diffusion_fMRI.

## Acknowledgments

We thank Dimitri Van De Ville, Essa Yacoub, Eleonora Fornari, Lijing Xin, Olivier Reynaud and Jelle Veraart for much valuable advice and very helpful discussions. This work was supported by the Swiss National Science Foundation under a Spark award CRSK-2_190882 (to I.O.J.). We are grateful for the provision of simultaneous multi-slice (multiband) pulse sequence and reconstruction algorithms from the Center for Magnetic Resonance Research, University of Minnesota. We acknowledge access to the facilities and expertise of the CIBM Center for Biomedical Imaging, a Swiss research center of excellence founded and supported by Lausanne University Hospital (CHUV), University of Lausanne (UNIL), Ecole Polytechnique Fédérale de Lausanne (EPFL), University of Geneva (UNIGE) and Geneva University Hospitals (HUG).

## Supplementary Data

**Fig. S1.**
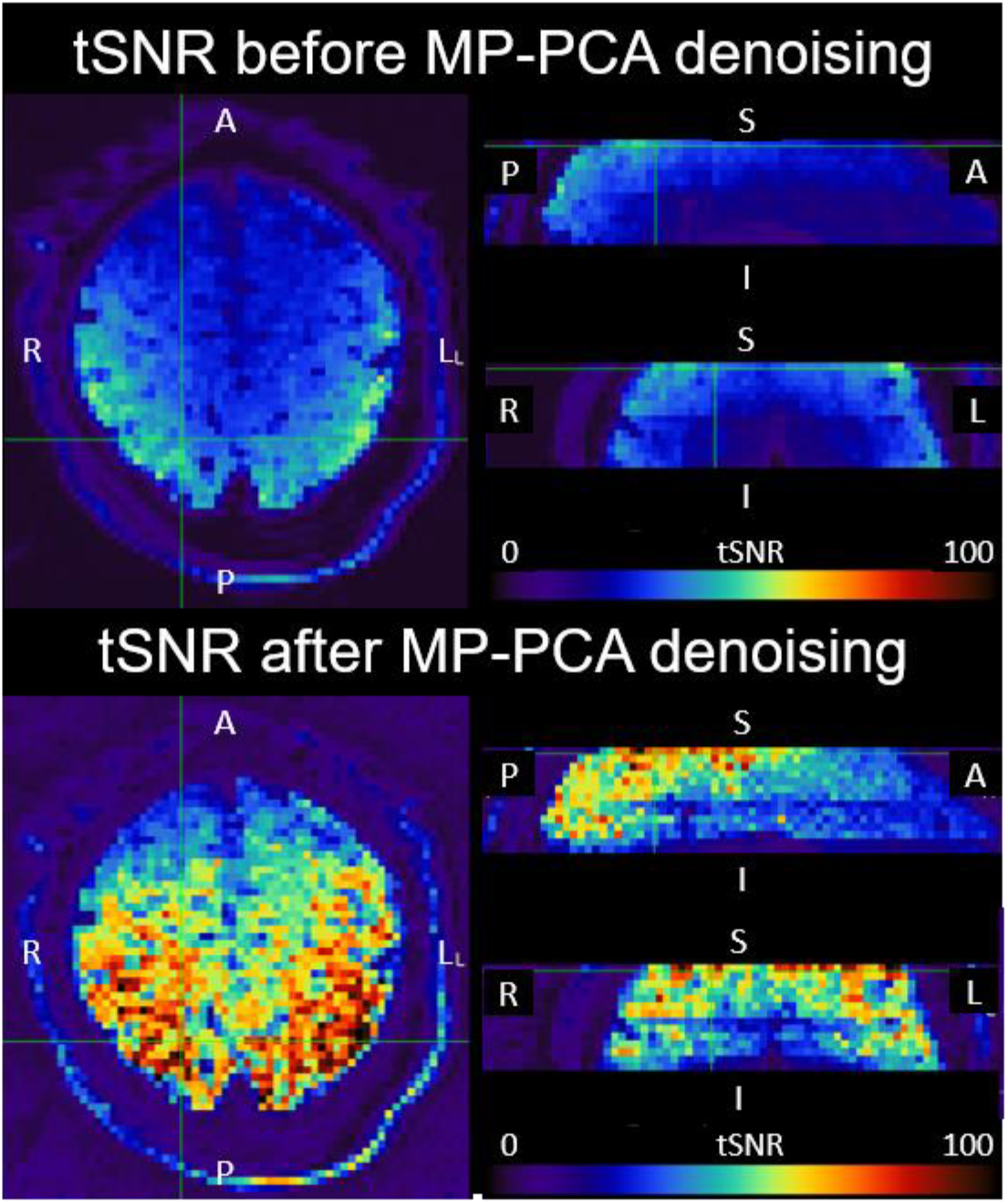
Temporal SNR maps before (top panel) and after (bottom panel) MP-PCA denoising for *b*=0.2 volumes of an exemplary subject. The denoising improved tSNR by a factor of almost two.

**Fig. S2.**
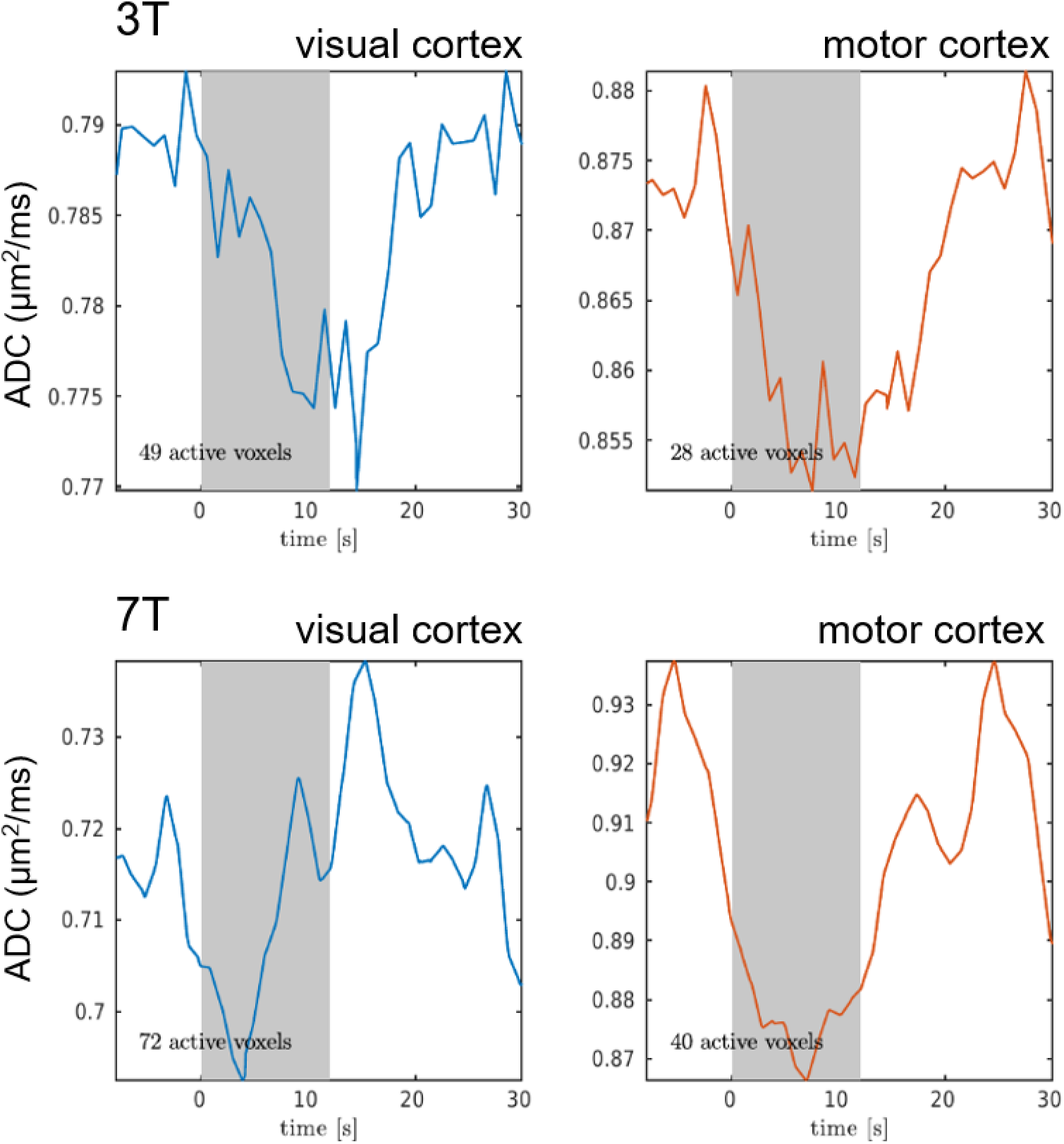
Average ADC responses in active voxels (z-statistic > 3.1) in the visual and motor cortices for an individual subject, at 3T (top) and 7T (bottom). The individual response characteristics mirror the group-level analysis and demonstrate a dfMRI response can be recorded at the single-subject level in physiological conditions.

**Fig. S3.**
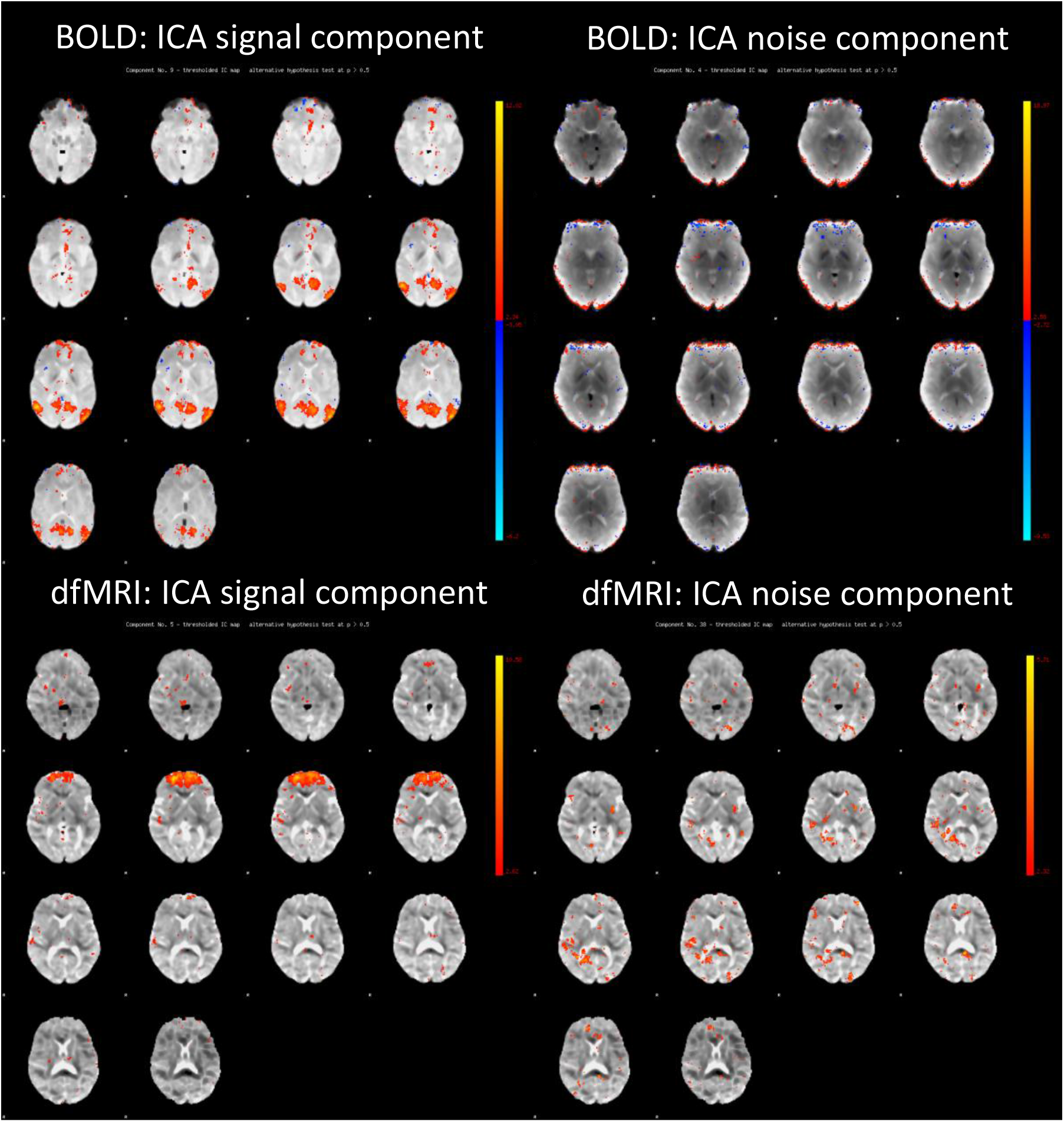
Exemplary signal and noise ICA components in SE BOLD and ADC-dfMRI datasets.

**Fig. S4.**
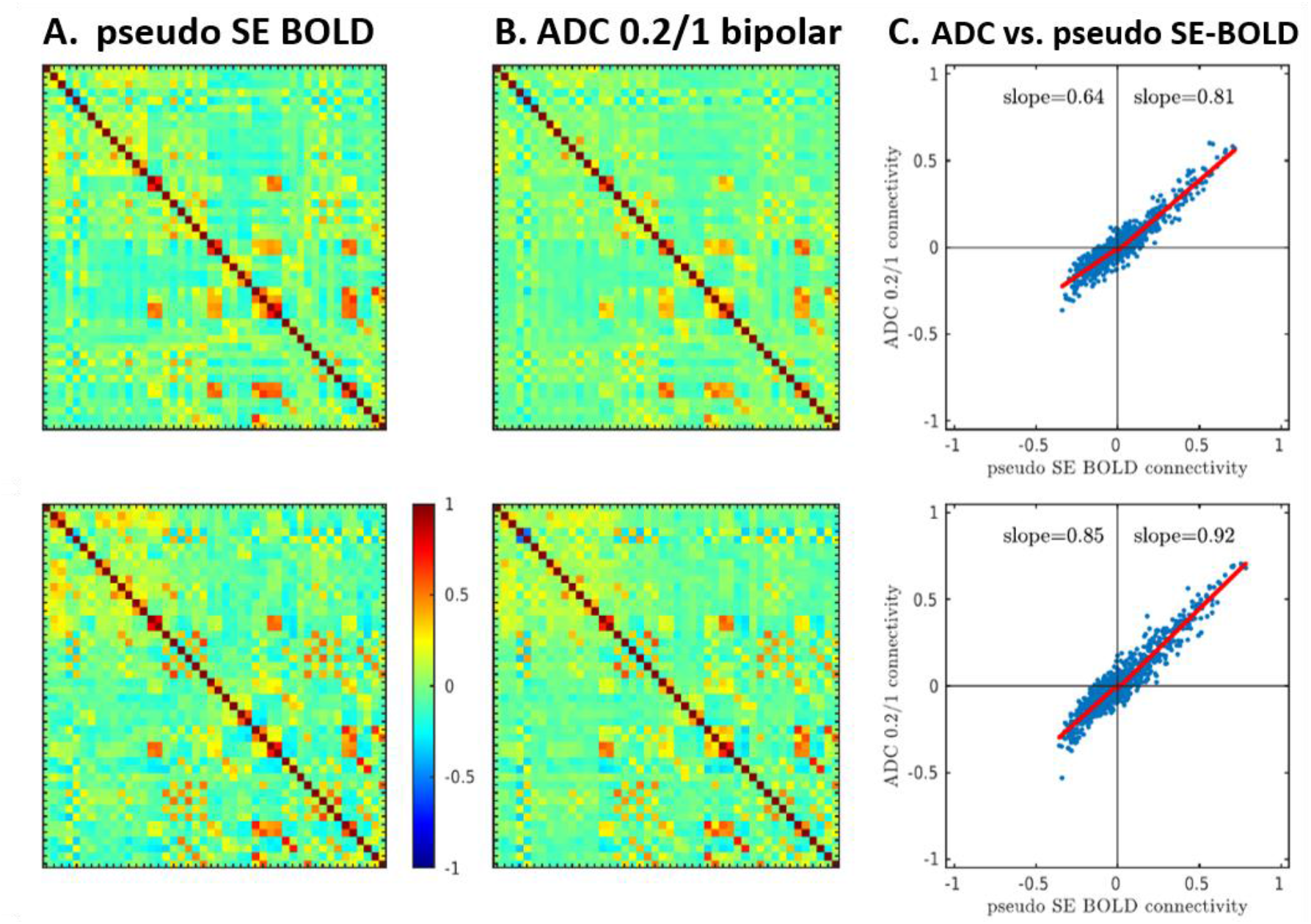
Resting-state analysis following manual ICA cleaning and GSR. Group averages of FC matrices from pseudo SE BOLD (*b*=0.2 timecourses) (**A**) and ADC *b*=0.2/1 (**B**) at 3T and 7T. **C**: Correlation of FC strength derived from pseudo SE BOLD vs. dfMRI. FC coefficients derived from both methods agree more strongly than BOLD vs. dfMRI (**Fig. 5**), including negative correlations.

**Fig. S5.**
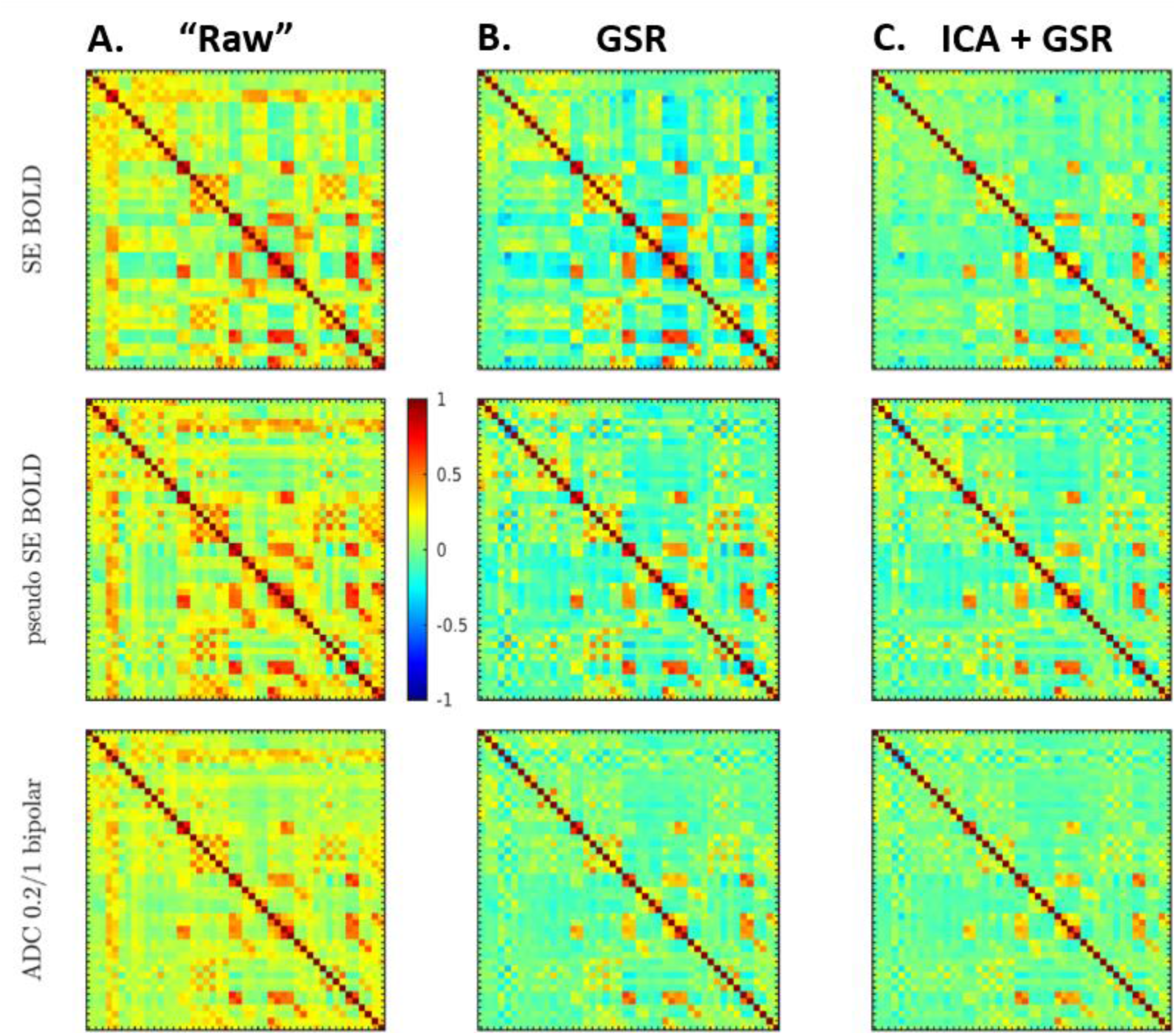
Average functional connectivity matrices at 3T, derived from either “raw” signal (**A**), after GSR (**B**) or after manual ICA cleaning followed by GSR (**C**). Rows correspond to RS-FC derived from various protocols: SE BOLD, pseudo SE BOLD (*b*=0.2 time-courses) and ADC (from pairs *b*=0.2/1). While ICA cleaning affects SE BOLD connectivity – by removing unwanted physiological artifacts, it has little impact on ADC-derived connectivity. By intrinsically removing large field fluctuation contributions to functional contrast, such as those of vascular origin, ADC dfMRI also has a valuable attribute of yielding data that is “*clean*” by design.

**Fig. S6.**
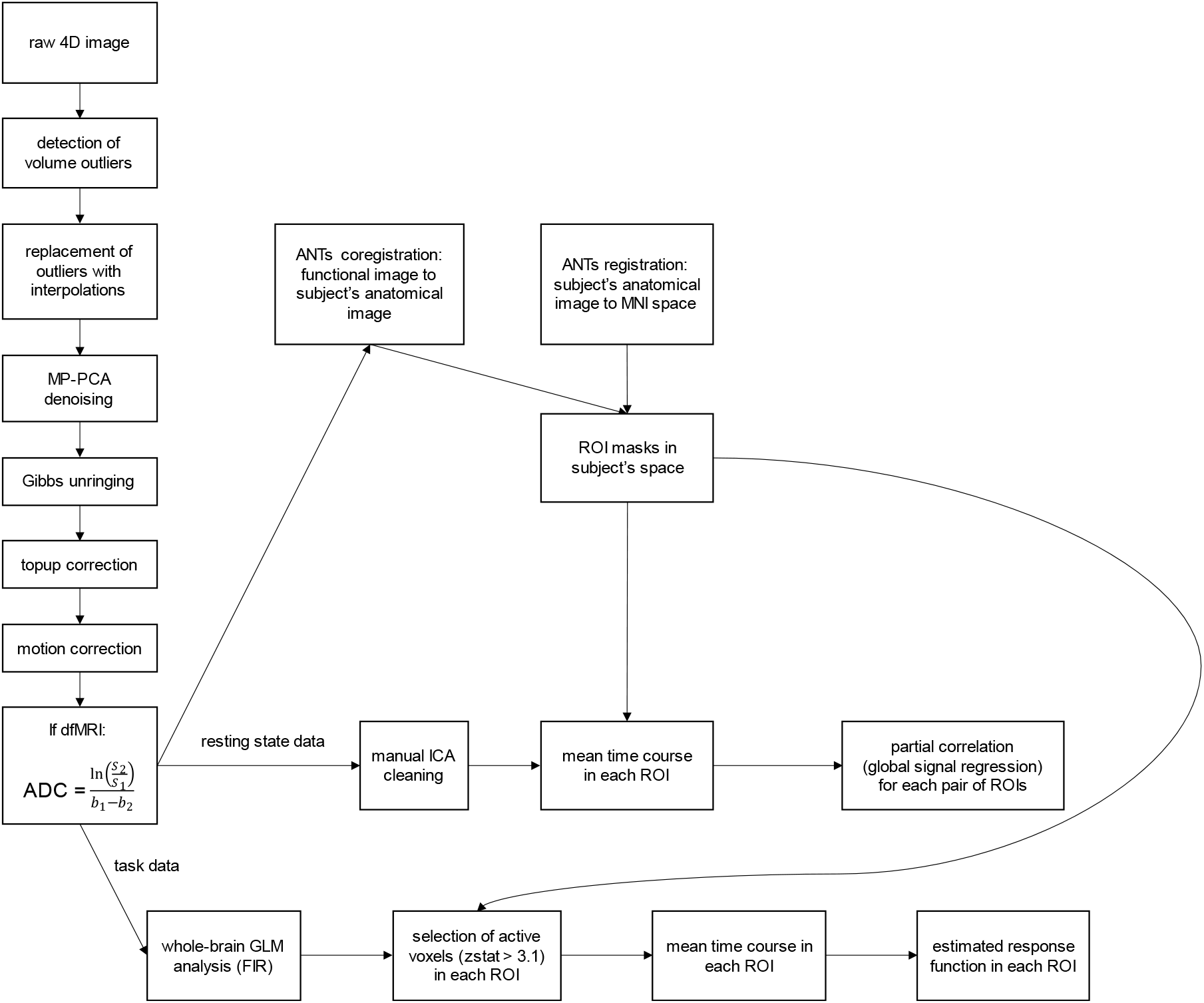
The employed processing pipeline. Data pre-processing included denoising (1, 2), Gibbs unringing (3) and corrections for susceptibility distortion (4) and motion. For the apparent diffusion coefficient (ADC) calculation, *S_i_* is the signal level measured with diffusion-weighting *b_i_*. All the processing scripts are at https://github.com/wiktorolszowy/diffusion_fMRI.

**Fig. S7.**
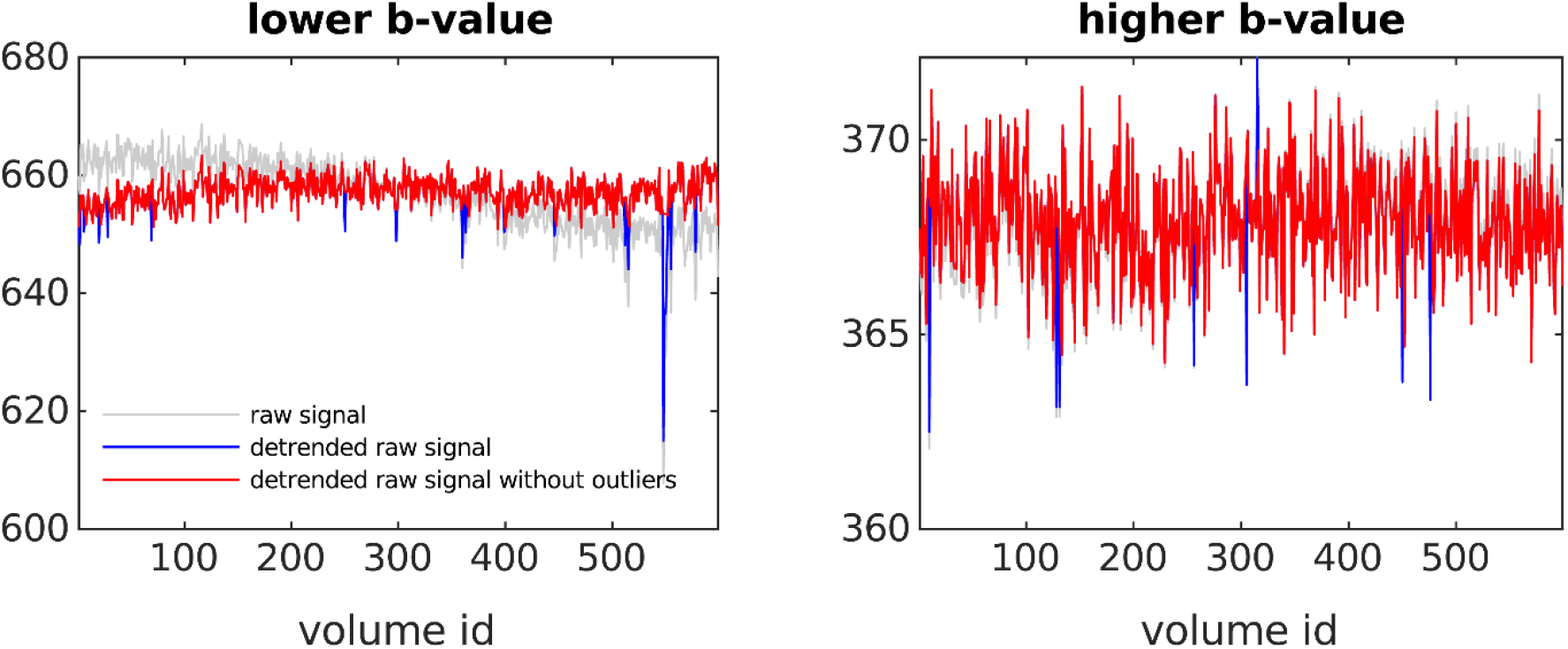
Outlier detection and removal for an exemplary dataset. For each volume, the mean across the volume was calculated (gray line). As low-frequency drifts could affect the detection of volume outliers, the signal was then linearly detrended (blue line). A volume was considered an outlier if its detrended average signal was away from the median across all the volumes by more than a pre-specified fraction of this median. This threshold fraction was set at 1% and 3% for 3T and 7T, respectively. Finally, the signal for outliers was replaced with linear interpolation (red line).

**Table S1.** Acquisition parameters for the four fMRI protocols compared.

| | | (1) SE BOLD | | (2) ADC from $b=0.2/1$, bipolar | | (3) ADC from $b=0/1$, bipolar | | (4) ADC from $b=0.2/1$, monopolar | |
| --- | --- | --- | --- | --- | --- | --- | --- | --- | --- |
|  |  | 3T | 7T | 3T | 7T | 3T | 7T | 3T | 7T |
| Task | # Datasets | 5 | 8 | 19 | 13 | 1 | 2 | 8 | 8 |
|  | TE (ms) | 73 | 65 | 72 | 62 | 72 | 65 | 72 | 62 |
|  | TR (s) | 1 |  |  |  |  |  |  |  |
|  | Matrix | 94x94 |  |  |  |  | 116x116 | 94x94 |  |
|  | Resolution (mm<sup>2</sup>) | 2.5x2.5 |  |  |  |  | 2x2 | 2.5x2.5 |  |
|  | Slices | 16 |  |  |  |  |  |  |  |
|  | Thickness (mm) | 2.5 |  |  |  |  | 2 | 2.5 |  |
|  | Total volumes | 600 |  |  |  |  |  |  |  |
| Resting-state | # Datasets | 10 | 10 | 12 | 11 | 0 | 2 | 5 | 5 |
|  | TE (ms) | 73 | 65 | 73 | 65 |  | 65 | 73 | 65 |
|  | TR (s) | 1 |  |  |  |  |  |  |  |
|  | Matrix | 116x116 |  |  |  |  |  |  |  |
|  | Resolution (mm<sup>2</sup>) | 2x2 |  |  |  |  |  |  |  |
|  | Slices | 16 |  |  |  |  |  |  |  |
|  | Thickness (mm) | 2 |  |  |  |  |  |  |  |
|  | Total volumes | 600 |  |  |  |  |  |  |  |

**Table S2.**
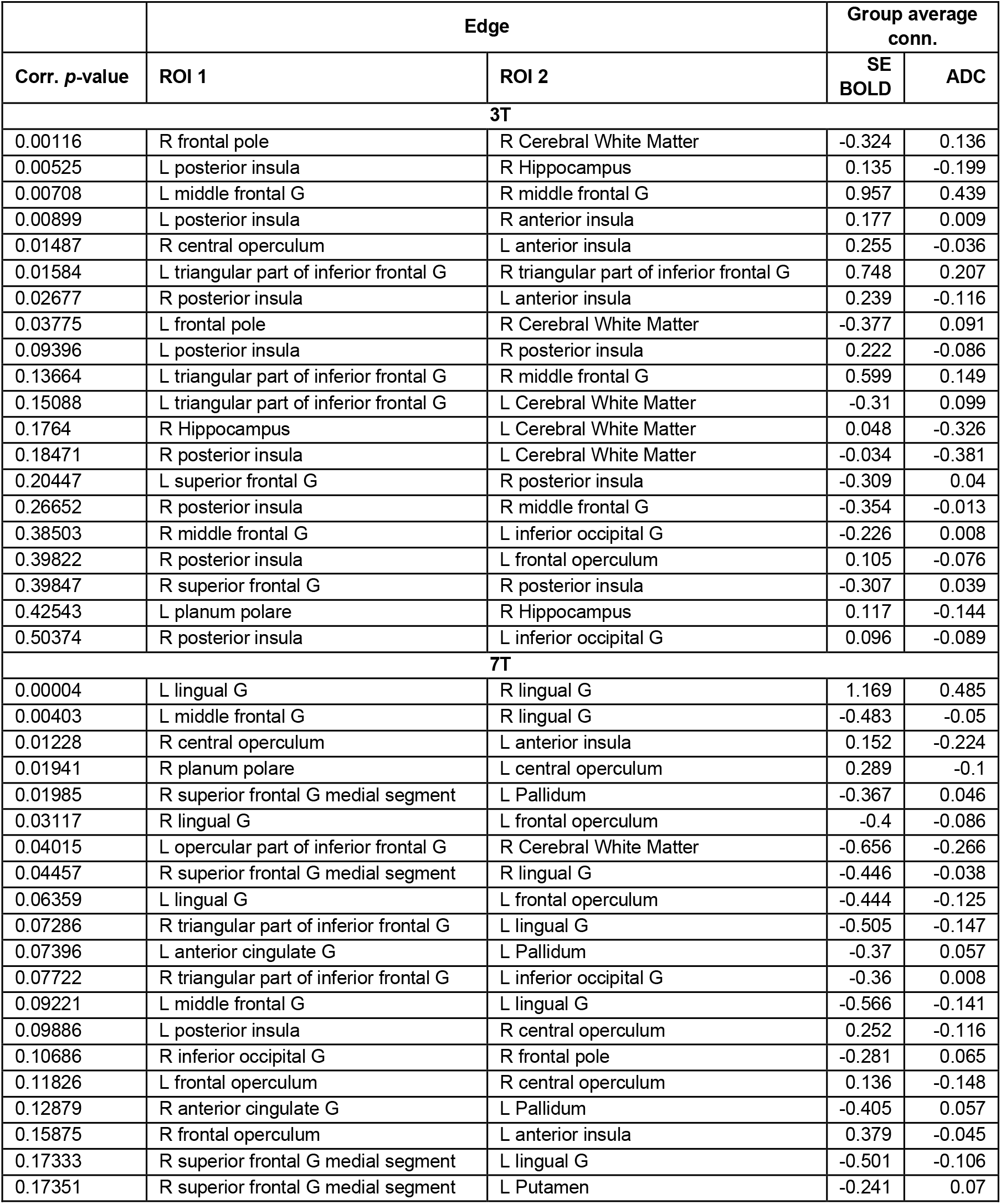
Resting-state functional connectivity edges with most significant differences between SE BOLD and ADC *b*=0.2/1 bipolar, dependent on the field strength, following GSR without manual ICA cleaning. ROIs come from the Neuromorphometrics atlas. The edges are sorted after Bonferroni-corrected *p*-values. The *p*-values come from two-sample *t*-tests on Fisher-transformed Pearson correlation coefficients. The last two columns show group averages of the Fisher-transformed Pearson correlation coefficients. The majority of edges with significantly different connectivity had either a low or negative correlation in BOLD, which was further attenuated in dfMRI. R/L: Right/Left; G: Gyrus.

| Corr. $p$-value | Edge | | Group average conn. | |
| --- | --- | --- | --- | --- |
|  | ROI 1 | ROI 2 | SE BOLD | ADC |
| **3T** |  |  |  |  |
| 0.00116 | R frontal pole | R Cerebral White Matter | -0.324 | 0.136 |
| 0.00525 | L posterior insula | R Hippocampus | 0.135 | -0.199 |
| 0.00708 | L middle frontal G | R middle frontal G | 0.957 | 0.439 |
| 0.00899 | L posterior insula | R anterior insula | 0.177 | 0.009 |
| 0.01487 | R central operculum | L anterior insula | 0.255 | -0.036 |
| 0.01584 | L triangular part of inferior frontal G | R triangular part of inferior frontal G | 0.748 | 0.207 |
| 0.02677 | R posterior insula | L anterior insula | 0.239 | -0.116 |
| 0.03775 | L frontal pole | R Cerebral White Matter | -0.377 | 0.091 |
| 0.09396 | L posterior insula | R posterior insula | 0.222 | -0.086 |
| 0.13664 | L triangular part of inferior frontal G | R middle frontal G | 0.599 | 0.149 |
| 0.15088 | L triangular part of inferior frontal G | L Cerebral White Matter | -0.31 | 0.099 |
| 0.1764 | R Hippocampus | L Cerebral White Matter | 0.048 | -0.326 |
| 0.18471 | R posterior insula | L Cerebral White Matter | -0.034 | -0.381 |
| 0.20447 | L superior frontal G | R posterior insula | -0.309 | 0.04 |
| 0.26652 | R posterior insula | R middle frontal G | -0.354 | -0.013 |
| 0.38503 | R middle frontal G | L inferior occipital G | -0.226 | 0.008 |
| 0.39822 | R posterior insula | L frontal operculum | 0.105 | -0.076 |
| 0.39847 | R superior frontal G | R posterior insula | -0.307 | 0.039 |
| 0.42543 | L planum polare | R Hippocampus | 0.117 | -0.144 |
| 0.50374 | R posterior insula | L inferior occipital G | 0.096 | -0.089 |
| **7T** |  |  |  |  |
| 0.00004 | L lingual G | R lingual G | 1.169 | 0.485 |
| 0.00403 | L middle frontal G | R lingual G | -0.483 | -0.05 |
| 0.01228 | R central operculum | L anterior insula | 0.152 | -0.224 |
| 0.01941 | R planum polare | L central operculum | 0.289 | -0.1 |
| 0.01985 | R superior frontal G medial segment | L Pallidum | -0.367 | 0.046 |
| 0.03117 | R lingual G | L frontal operculum | -0.4 | -0.086 |
| 0.04015 | L opercular part of inferior frontal G | R Cerebral White Matter | -0.656 | -0.266 |
| 0.04457 | R superior frontal G medial segment | R lingual G | -0.446 | -0.038 |
| 0.06359 | L lingual G | L frontal operculum | -0.444 | -0.125 |
| 0.07286 | R triangular part of inferior frontal G | L lingual G | -0.505 | -0.147 |
| 0.07396 | L anterior cingulate G | L Pallidum | -0.37 | 0.057 |
| 0.07722 | R triangular part of inferior frontal G | L inferior occipital G | -0.36 | 0.008 |
| 0.09221 | L middle frontal G | L lingual G | -0.566 | -0.141 |
| 0.09886 | L posterior insula | R central operculum | 0.252 | -0.116 |
| 0.10686 | R inferior occipital G | R frontal pole | -0.281 | 0.065 |
| 0.11826 | L frontal operculum | R central operculum | 0.136 | -0.148 |
| 0.12879 | R anterior cingulate G | L Pallidum | -0.405 | 0.057 |
| 0.15875 | R frontal operculum | L anterior insula | 0.379 | -0.045 |
| 0.17333 | R superior frontal G medial segment | L lingual G | -0.501 | -0.106 |
| 0.17351 | R superior frontal G medial segment | L Putamen | -0.241 | 0.07 |

## References

1. S. Ogawa, et al., Intrinsic signal changes accompanying sensory stimulation: functional brain mapping with magnetic resonance imaging. Proceedings of the National Academy of Sciences of the United States of America 89, 5951–5 (1992).

2. R. Turner, Uses, misuses, new uses and fundamental limitations of magnetic resonance imaging in cognitive science. Philosophical Transactions of the Royal Society B: Biological Sciences 371, 20150349 (2016).

3. A. Shmuel, E. Yacoub, D. Chaimow, N. K. Logothetis, K. Ugurbil, Spatio-temporal point-spread function of fMRI signal in human gray matter at 7 Tesla. NeuroImage 35, 539–552 (2007).

4. K. L. West, et al., BOLD hemodynamic response function changes significantly with healthy aging. NeuroImage 188, 198–207 (2019).

5. R. W. Pak, et al., Implications of neurovascular uncoupling in functional magnetic resonance imaging (fMRI) of brain tumors. J Cereb Blood Flow Metab 37, 3475–3487 (2017).

6. C. Mg, R. C, Vascular Disease in Patients with Multiple Sclerosis: A Review. J Vasc Med Surg 04 (2016).

7. J. C. Gore, et al., Functional MRI and resting state connectivity in white matter - a mini-review. Magnetic Resonance Imaging 63, 1–11 (2019).

8. L. A. Grajauskas, T. Frizzell, X. Song, R. C. N. D’Arcy, White Matter fMRI Activation Cannot Be Treated as a Nuisance Regressor: Overcoming a Historical Blind Spot. Front. Neurosci. 13 (2019).

9. J. Dubois, R. Adolphs, Building a Science of Individual Differences from fMRI. Trends in Cognitive Sciences 20, 425–443 (2016).

10. D. Le Bihan, S. Urayama, T. Aso, T. Hanakawa, H. Fukuyama, Direct and fast detection of neuronal activation in the human brain with diffusion MRI. Proceedings of the National Academy of Sciences 103, 8263 (2006).

11. R. D. Andrew, B. A. Macvicar, Imaging cell volume changes and neuronal excitation in the hippocampal slice. Neuroscience 62, 371–383 (1994).

12. F. Lang, et al., Functional Significance of Cell Volume Regulatory Mechanisms. Physiological Reviews 78, 247–306 (1998).

13. A. D. Sherpa, F. Xiao, N. Joseph, C. Aoki, S. Hrabetova, Activation of beta-adrenergic receptors in rat visual cortex expands astrocytic processes and reduces extracellular space volume. Synapse (New York, N.Y.) 70, 307–16 (2016).

14. A. Darquie, J. B. Poline, C. Poupon, H. Saint-Jalmes, D. Le Bihan, Transient decrease in water diffusion observed in human occipital cortex during visual stimulation. Proceedings of the National Academy of Sciences of the United States of America 98, 9391–5 (2001).

15. T. Tsurugizawa, L. Ciobanu, D. Le Bihan, Water diffusion in brain cortex closely tracks underlying neuronal activity. Proceedings of the National Academy of Sciences 110, 11636 (2013).

16. D. Nunes, A. Ianus, N. Shemesh, Layer-specific connectivity revealed by diffusion-weighted functional MRI in the rat thalamocortical pathway. NeuroImage 184, 646–657 (2019).

17. J. Flint, B. Hansen, P. Vestergaard-Poulsen, S. J. Blackband, Diffusion weighted magnetic resonance imaging of neuronal activity in the hippocampal slice model. NeuroImage 46, 411–8 (2009).

18. N. Tirosh, U. Nevo, Neuronal activity significantly reduces water displacement: DWI of a vital rat spinal cord with no hemodynamic effect. NeuroImage 76, 98–107 (2013).

19. Y. Abe, T. Tsurugizawa, D. Le Bihan, Water diffusion closely reveals neural activity status in rat brain loci affected by anesthesia. PLOS Biology 15, e2001494 (2017).

20. K. L. Miller, et al., Evidence for a vascular contribution to diffusion FMRI at high b value. Proceedings of the National Academy of Sciences 104, 20967–20972 (2007).

21. T. Jin, S.-G. Kim, Functional changes of apparent diffusion coefficient during visual stimulation investigated by diffusion-weighted gradient-echo fMRI. NeuroImage 41, 801–812 (2008).

22. J. A. A. Autio, et al., High b-value diffusion-weighted fMRI in a rat forepaw electrostimulation model at 7 T. NeuroImage 57, 140–148 (2011).

23. R. Bai, C. V. Stewart, D. Plenz, P. J. Basser, Assessing the sensitivity of diffusion MRI to detect neuronal activity directly. Proc Natl Acad Sci U S A 113, E1728–E1737 (2016).

24. A. D. Luca, L. Schlaffke, J. C. W. Siero, M. Froeling, A. Leemans, On the sensitivity of the diffusion MRI signal to brain activity in response to a motor cortex paradigm. Human Brain Mapping 40, 5069–5082 (2019).

25. A. Pampel, T. H. Jochimsen, H. E. Möller, BOLD background gradient contributions in diffusion-weighted fMRI-comparison of spin-echo and twice-refocused spin-echo sequences. NMR Biomed. 23, 610–618 (2010).

26. W. M. Spees, et al., MRI-based assessment of function and dysfunction in myelinated axons. Proceedings of the National Academy of Sciences of the United States of America 115, E10225–E10234 (2018).

27. D. Nunes, R. Gil, N. Shemesh, A rapid-onset diffusion functional MRI signal reflects neuromorphological coupling dynamics. NeuroImage, 117862 (2021).

28. B. Ades-Aron, et al., Improved Task-based Functional MRI Language Mapping in Patients with Brain Tumors through Marchenko-Pastur Principal Component Analysis Denoising. Radiology 298, 365–373 (2021).

29. Y. Diao, T. Yin, R. Gruetter, I. O. Jelescu, PIRACY: An Optimized Pipeline for Functional Connectivity Analysis in the Rat Brain. Front. Neurosci. 15, 602170 (2021).

30. J. Veraart, et al., Denoising of diffusion MRI using random matrix theory. NeuroImage 142, 394–406 (2016).

31. G. H. Glover, Deconvolution of Impulse Response in Event-Related BOLD fMRI. NeuroImage 9, 416–429 (1999).

32. M. J. Sætra, G. T. Einevoll, G. Halnes, An electrodiffusive neuron-extracellular-glia model with somatodendritic interactions. bioRxiv, 2020.07.13.200287 (2020).

33. L. Huber, et al., High-Resolution CBV-fMRI Allows Mapping of Laminar Activity and Connectivity of Cortical Input and Output in Human M1. Neuron 96, 1253–1263.e7 (2017).

34. S. Mori, Ed., MRI atlas of human white matter, 1. ed (Elsevier, 2005).

35. J. R. Gawryluk, E. L. Mazerolle, R. C. N. D’Arcy, Does functional MRI detect activation in white matter? A review of emerging evidence, issues, and future directions. Front. Neurosci. 8 (2014).

36. W. M. Spees, T.-H. Lin, S.-K. Song, White-matter diffusion fMRI of mouse optic nerve. NeuroImage 65, 209–215 (2013).

37. T.-H. Lin, et al., Diffusion fMRI detects white-matter dysfunction in mice with acute optic neuritis. Neurobiology of Disease 67, 1–8 (2014).

38. J. Li, et al., Global signal regression strengthens association between resting-state functional connectivity and behavior. NeuroImage 196, 126–141 (2019).

39. K. Murphy, R. M. Birn, D. A. Handwerker, T. B. Jones, P. A. Bandettini, The impact of global signal regression on resting state correlations: Are anti-correlated networks introduced? NeuroImage 44, 893–905 (2009).

40. M. D. Fox, et al., The human brain is intrinsically organized into dynamic, anticorrelated functional networks. Proceedings of the National Academy of Sciences of the United States of America 102, 9673 (2005).

41. A. Nalci, B. D. Rao, T. T. Liu, Global signal regression acts as a temporal downweighting process in resting-state fMRI. NeuroImage 152, 602–618 (2017).

42. M. D. Fox, D. Zhang, A. Z. Snyder, M. E. Raichle, The global signal and observed anticorrelated resting state brain networks. Journal of neurophysiology 101, 3270–83 (2009).

43. M. Bianciardi, M. Fukunaga, P. van Gelderen, J. A. de Zwart, J. H. Duyn, Negative BOLD-fMRI Signals in Large Cerebral Veins. J Cereb Blood Flow Metab 31, 401–412 (2011).

44. A. Devor, et al., Suppressed Neuronal Activity and Concurrent Arteriolar Vasoconstriction May Explain Negative Blood Oxygenation Level-Dependent Signal. Journal of Neuroscience 27, 4452–4459 (2007).

45. Y. Abe, et al., Diffusion functional MRI reveals global brain network functional abnormalities driven by targeted local activity in a neuropsychiatric disease mouse model. NeuroImage 223, 117318 (2020).

46. Jelescu, I.O., Resting-state diffusion fMRI bears strong resemblance and only subtle differences to BOLD fMRI. ISMRM (2019).

47. J. A. Fraser, C. L.-H. Huang, A quantitative analysis of cell volume and resting potential determination and regulation in excitable cells. The Journal of Physiology 559, 459–478 (2004).

48. A. E. Campbell-Washburn, et al., Opportunities in Interventional and Diagnostic Imaging by Using High-Performance Low-Field-Strength MRI. Radiology 293, 384–393 (2019).

49. Y. Wang, P. Gelderen, J. A. Zwart, A. E. Campbell-Washburn, J. H. Duyn, FMRI based on transition-band balanced SSFP in comparison with EPI on a high-performance 0.55 T scanner. Magn Reson Med 85, 3196–3210 (2021).

50. T. Jin, S.-G. Kim, Change of the cerebrospinal fluid volume during brain activation investigated by T1ρ-weighted fMRI. NeuroImage 51, 1378–1383 (2010).

51. B. P. Thomas, et al., Physiologic underpinnings of negative BOLD cerebrovascular reactivity in brain ventricles. NeuroImage 83, 505–512 (2013).

52. M.-C. Chou, et al., FLAIR Diffusion-Tensor MR Tractography: Comparison of Fiber Tracking with Conventional Imaging. 7 (2005).

53. T. Aso, et al., An intrinsic diffusion response function for analyzing diffusion functional MRI time series. NeuroImage 47, 1487–1495 (2009).

54. W. Olszowy, J. Aston, C. Rua, G. B. Williams, Accurate autocorrelation modeling substantially improves fMRI reliability. Nat Commun 10, 1220 (2019).

55. K. Uğurbil, et al., Pushing spatial and temporal resolution for functional and diffusion MRI in the Human Connectome Project. NeuroImage 80, 80–104 (2013).

56. M. E. Raichle, et al., A default mode of brain function. Proceedings of the National Academy of Sciences 98, 676–682 (2001).

57. J. L. R. Andersson, S. Skare, J. Ashburner, How to correct susceptibility distortions in spin-echo echo-planar images: application to diffusion tensor imaging. NeuroImage 20, 870–888 (2003).

58. J. Peirce, et al., PsychoPy2: Experiments in behavior made easy. Behav Res 51, 195–203 (2019).

59. E. Kellner, B. Dhital, V. G. Kiselev, M. Reisert, Gibbs-ringing artifact removal based on local subvoxel-shifts: Gibbs-Ringing Artifact Removal. Magn. Reson. Med. 76, 1574–1581 (2016).

60. F. Alfaro-Almagro, et al., Image processing and Quality Control for the first 10,000 brain imaging datasets from UK Biobank. NeuroImage 166, 400–424 (2018).

61. Karl. J. Friston, et al., Spatial registration and normalization of images. Hum. Brain Mapp. 3, 165–189 (1995).

62. K. J. Friston, Ed., Statistical parametric mapping: the analysis of funtional brain images, 1st ed (Elsevier/Academic Press, 2007).

63. B. B. Avants, et al., A reproducible evaluation of ANTs similarity metric performance in brain image registration. NeuroImage 54, 2033–2044 (2011).

64. B. B. Avants, et al., The optimal template effect in hippocampus studies of diseased populations. NeuroImage 49, 2457–2466 (2010).

65. J. R. Polimeni, V. Renvall, N. Zaretskaya, B. Fischl, Analysis strategies for high-resolution UHF-fMRI data. NeuroImage 168, 296–320 (2018).

66. M. Jenkinson, C. F. Beckmann, T. E. J. Behrens, M. W. Woolrich, S. M. Smith, FSL. NeuroImage 62, 782–790 (2012).

67. R. Henson, M. D. Rugg, K. J. Friston, The choice of basis functions in event-related fMRI. NeuroImage 13, 149 (2001).

68. C. F. Beckmann, S. M. Smith, Probabilistic Independent Component Analysis for Functional Magnetic Resonance Imaging. IEEE Trans. Med. Imaging 23, 137–152 (2004).

69. L. Griffanti, et al., Hand classification of fMRI ICA noise components. NeuroImage 154, 188–205 (2017).

70. K. L. Miller, P. Jezzard, Modeling SSFP functional MRI contrast in the brain. Magn. Reson. Med. 60, 661–673 (2008).

## SI References

1. J. Veraart, et al., Denoising of diffusion MRI using random matrix theory. NeuroImage 142, 394–406 (2016).

2. B. Ades-Aron, et al., Improved Task-based Functional MRI Language Mapping in Patients with Brain Tumors through Marchenko-Pastur Principal Component Analysis Denoising. Radiology 298, 365–373 (2021).

3. E. Kellner, B. Dhital, V. G. Kiselev, M. Reisert, Gibbs-ringing artifact removal based on local subvoxel-shifts: Gibbs-Ringing Artifact Removal. Magn. Reson. Med. 76, 1574–1581 (2016).

4. J. L. R. Andersson, S. Skare, J. Ashburner, How to correct susceptibility distortions in spin-echo echo-planar images: application to diffusion tensor imaging. NeuroImage 20, 870–888 (2003).

